# Upstream open reading frame translation enhances immunogenic peptide presentation in mitotically arrested cancer cells

**DOI:** 10.1101/2025.01.08.631915

**Authors:** Alexander Kowar, Jonas P. Becker, Rossella Del Pizzo, Zhiwei Tang, Julien Champagne, Pierre-René Körner, Jasmine Montenegro Navarro, Fiona Megan Tilghman, Hanan Sakeer, Angelika B. Riemer, Reuven Agami, Fabricio Loayza-Puch

## Abstract

Mitosis is a critical phase of the cell cycle and a vulnerable point where cancer cells can be effectively disrupted, leading to cell death and inhibition of tumor growth. However, challenges such as drug resistance remain significant in clinical applications. During mitosis, mRNA translation is generally downregulated, while non-canonical translation of specific transcripts proceeds. Here, we demonstrate that mitotic cancer cells redistribute ribosomes toward the 5’ untranslated region (5’ UTR) and the start of the coding sequence (CDS), enhancing the translation of thousands of upstream open reading frames (uORFs) and upstream overlapping open reading frames (uoORFs). This mitotic induction of uORF/uoORF enriches the presentation of immunopeptides at the surface of cancer cells following treatment with mitotic inhibitors. Functional assays indicate the potential of such neoepitopes to provoke cancer-cell killing by T cells. Altogether, our findings highlight the therapeutic potential of targeting uORF/uoORF-derived neoepitopes in combination with mitotic inhibitors to enhance immune recognition and tumor cell elimination.

## Introduction

Mitosis is a critical phase of cellular reproduction and a key target for cancer therapy^1^. Although it is typically the shortest stage of the mammalian cell cycle, mitosis involves profound changes in cellular organization. The control of translation during mitosis plays an essential role in regulating the cell cycle, with particular significance in cancer biology. Most mRNAs undergo gene-specific translational downregulation during mitosis, rather than activation^2,3^. However, transcripts with a terminal oligopyrimidine (TOP) tract can escape this global translational suppression^4^. Cyclin-dependent kinase 1 (CDK1) plays an important role in regulating mRNA translation during the M-phase of the cell cycle. Recent studies show that CDK1 influences translational regulation by phosphorylating specific substrates involved in mRNA translation. For example, CDK1 can phosphorylate ribosomal proteins and translation initiation factors, enhancing the protein synthesis needed for mitotic progression^5–7^. This activity ensures the cell possesses the necessary components for successful division.

Since cancer cells frequently bypass internal mitotic checkpoints, they often present high sensitivity to mitotic inhibitors^8^. These agents disrupt the microtubule dynamics required for chromosome segregation during cell division, leading to mitotic arrest and, ultimately, cell death^9,10^. Paclitaxel (Taxol), a widely used chemotherapeutic drug, acts by stabilizing microtubules and preventing their depolymerization^11,12^. It binds to the β-subunit of tubulin, locking the microtubule structure in place and inhibiting the dynamic reorganization of the microtubule network. This stabilization causes cell cycle arrest in the G2/M phase, which ultimately triggers apoptosis^13,14^. Clinically, paclitaxel is employed as both a first-line and subsequent therapy for several advanced carcinomas, including ovarian, breast, lung, and pancreatic cancers^15^. Its unique mechanism of action makes it effective against a broad range of malignancies, though resistance development remains a significant challenge in cancer treatment.

Non-canonical translation has emerged as an important mechanism for remodeling the proteome and immunopeptidome, influencing cellular responses and immune recognition^16,17^. Non-canonical open reading frames (ncORFs) have gained attention due to advancements in ribosome profiling (RiboSeq), which have allowed for the identification of actively translated ncORFs encoded by long noncoding RNAs (lncRNAs), pseudogenes, and untranslated regions^18–23^. Recent studies have revealed thousands of ncORFs, some of which play critical biological roles, such as regulating cell proliferation and serving as neoantigens presented by major histocompatibility complex I (MHC I)^24,25^. Notably, emerging research has characterized several ncORF-derived microproteins with crucial functions in development and muscle biology^26–28^. However, the potential of ncORF-derived peptides to enhance the effectiveness of existing chemotherapies as immune targets remains largely unexplored.

In this study, we investigated the dynamics of noncanonical translation in cancer cells arrested in mitosis. Using ribosome profiling, we observed a significant shift of ribosomes toward the 5’ UTR, leading to an increased translation of uORF/uoORFs. Additionally, we identified several uORF/uoORF-derived peptides presented on the surface of cancer cells, suggesting their potential role in shaping the immune response and serving as tumor antigens. Upon mitotic arrest induced by chemotherapy, we observed an enhanced presentation of specific uORF/uoORF derived peptides, which were recognized by CD8^+^ T cells, thus promoting tumor cell killing. These findings underscore the potential of targeting therapy-induced uORF/uoORF-derived peptides as a novel strategy in cancer immunotherapy, offering new opportunities for cancer treatment.

## Results

### Extensive ribosome redistribution in cancer cells arrested in mitosis

To investigate the translational dynamics of cancer cells arrested in mitosis, we performed ribosome profiling on the osteosarcoma cell line U2OS, treated with Nocodazole—a drug that disrupts microtubule polymerization and halts cell cycle progression at mitosis (Fig. 1a). In proliferating cells, the global transcript distribution of ribosome protected fragments (RPFs) was uniform across the coding sequence (CDS). However, in mitotically arrested cells, there was a pronounced redistribution of ribosomes toward the 5’ UTR and the start of the CDS (Fig. 1b). The proportion of RPFs mapping to the 5’ UTR increased approximately two-fold compared to those mapping to the 3’ UTR (Fig. 1c). This redistribution was similarly observed across multiple cancer cell lines arrested in mitosis (Supplementary Fig. 1a-f).

**Figure 1.**
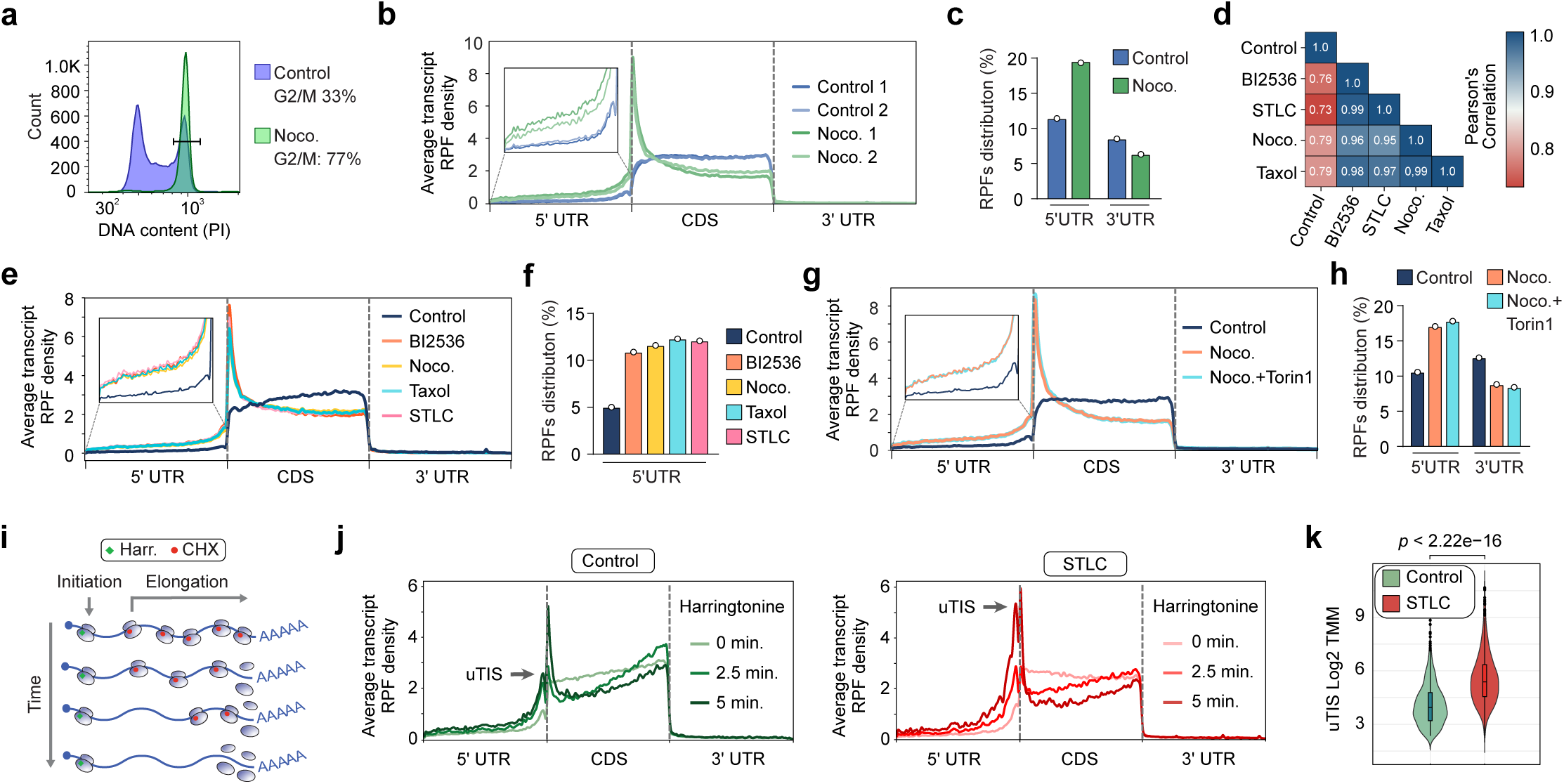
Prolonged mitotic arrest leads to ribosome redistribution toward the 5’ UTR. **a**, Representative propidium iodide staining of U2OS cells arrested in mitosis with Nocodazole (Noco. 0.5 µM) for 16 hours. **b,** Metagene profiles of ribosome-protected fragments (RPFs) in proliferating and mitotically arrested U2OS cells treated with Nocodazole (0.5 µM; 16 hours). For each transcript, raw RPF coverage was normalized to the sum of transcript coverage divided by its length. The normalized coverage for each transcript within the window was then summed across all selected transcripts. **c,** Quantification of RPF distribution in the 5’ UTR and 3’UTR of U2OS cells treated with vehicle (Control) or Nocodazole (0.5 µM) for 16 hours. **d,** Heatmap of Pearson correlation coefficients between gene RPKM in U2OS cells treated with vehicle (Control), BI2536 (0.1 µM), Nocodazole (0.5 µM), Taxol (1 µM), or STLC (5 µM) for 16 hours. **e,** Metagene profiles of RPFs in U2OS cells treated with vehicle (Control), BI2536 (0.1 µM), Nocodazole (0.5 µM), Taxol (1 µM), or STLC (1 µM) for 16 hours. **f,** Quantification of RPF distribution in the 5’ UTR of cells described in e. **g,** Metagene profiles of RPFs in U2OS cells treated with vehicle (Control), Nocodazole (0.5 µM), or Nocodazole (0.5 µM) combined with Torin1 (250 nM). Cells were treated with Nocodazole for 16 hours and with Torin1 for 2 hours. **h,** Quantification of RPF distribution in the 5’ UTR and 3’UTR of cells described in g. **i,** Schematic of the run-off elongation experiment in U2OS cells. Harringtonine (2 µg/ml) was added at different time points, and cells were harvested in cold PBS (100 µg/ml CHX). CHX, Cycloheximide. **j,** Metagene profiles of RPFs in U2OS cells treated with vehicle (Control) or STLC (5 µM) for 16 hours. Cells were harvested as described in i. uTIS, upstream translation initiation site. **k**, Violin plots showing the trimmed mean of M-values (TMM) distribution of upstream translation initiation sites (uTIS) in U2OS cells treated with harringtonine as in i. Each point represents TMM at -20/+20 nucleotides from the predicted start site. The width of the violin indicates point density. The center line represents the median; upper and lower bounds represent the 75th and 25th percentiles, respectively. Whiskers represent minimum and maximum values. *p*-value by one-sided Fisher test, no correction for multiple testing.

Nocodazole-induced microtubule depolymerization can cause protein misfolding and aggregation^29^. These aggregates can activate the integrated stress response (ISR), leading to eIF2α phosphorylation, which inhibits global protein synthesis while increasing translation from unconventional 5’ start sites^30^. To test whether ribosome redistribution is linked to stress-induced eIF2α phosphorylation, we treated U2OS cells with the eIF2α phosphatase inhibitor salubrinal and performed ribosome profiling. Although salubrinal treatment resulted in transcriptional upregulation of stress-responsive genes (Supplementary Fig. 1g), it did not induce a ribosome shift toward the 5’UTR (Supplementary Fig. 1h-i). Moreover, to determine whether the shift in footprints is drug- or mitotic arrest-specific, we induced mitotic arrest in U2OS cells through various molecular mechanisms. We treated U2OS cells with BI-2536, a PLK1 inhibitor; S-Trityl-L-cysteine (STLC), an inhibitor of mitotic kinesin Eg5; or Taxol, a drug that stabilizes tubulin polymerization (Supplementary Fig. 1j). Translational activity was highly correlated across all mitotic arrest conditions (Fig. 1d). Ribosome profiling from all mitotic arrest treatments showed a similar extent of ribosome footprint redistribution and a concomitant increase in the proportion of RPFs in the 5’UTR (Fig. 1e-f), indicating that this redistribution is specific to mitotic arrest rather than a general stress response or drug-specific effect.

During mitosis, CDK1 substitutes for mTOR’s role in activating cap-dependent translation by phosphorylating key translational regulators like the translation initiation factor eIF4E-binding protein 1 (4E-BP1)^2,5,6,31^. To test whether ribosome redistribution during mitotic arrest is dependent on mTOR activity, we treated mitotic U2OS cells with Torin 1, a potent and selective inhibitor of the mTOR kinase. Ribosome profiling showed that mTOR inhibition during mitosis has no effect on ribosome redistribution nor the relative proportion of RPFs in the 5’UTR (Fig. 1g-h). Furthermore, we observed that mTOR inhibition does not prevent the hyperphosphorylation of mTOR targets, such as 4E-BP1, which is known to play a key role in translation initiation (Supplementary Fig. 1k). These findings indicate that mTOR is not the primary driver of ribosome redistribution during mitosis and that other mechanisms, such as CDK1-mediated phosphorylation of 4E-BP1, may be more critical in this process.

Next, to determine whether the redistribution of RPFs in cells arrested in mitosis is associated with increased initiation rates at unconventional sites within the 5’ UTR, we performed global run-off ribosome profiling analysis using harringtonine. Harringtonine is a compound that specifically inhibits translation initiation by binding to the ribosomal machinery and preventing the formation of the first peptide bond. This results in the accumulation of ribosomes at translation start sites while depleting elongating ribosomes^32,33^ (Fig. 1i). In proliferating U2OS cells, most translation initiation sites (TISs) that accumulated over time were located at the annotated open reading frames (ORFs). In contrast, mitotically arrested U2OS cells showed an increase in initiation events at upstream translation initiation sites (uTISs), which levels were comparable to those at the annotated ORFs (Fig. 1j-k). A similar increase in 5’ UTR initiation events was observed in MDA-MB-231 cells arrested in mitosis (Supplementary Fig. 1l). Altogether, our data suggest that prolonged mitotic arrest results in ribosome redistribution, leading to increased unconventional initiation rates at the 5’ UTR.

### Enhanced translation of uORF/uoORFs in mitotically arrested cancer cells

To systematically define the initiation events occurring in the 5’UTR of cells arrested in mitosis, we employed PRICE (Probabilistic Inference of Codon Activities by an EM Algorithm), a computational method specifically designed for identifying non-canonical ORFs from ribosome profiling data^34^ (Fig. 2a). After filtering out canonical coding sequences and truncated ORFs, we identified 1,444 distinct actively translated non-canonical ORFs in proliferating cells and over 2,600 in mitotically arrested U2OS cells treated with various agents (Supplementary Datasets 1-5). Notably, the proportion of actively translated uORFs and uoORFs more than doubled in mitotically arrested cells, regardless of the molecular mechanism inducing the arrest, while the proportion of other non-canonical ORFs remained stable (Fig. 2b). This phenomenon was corroborated by similar observations in PC3 and MDA-MB-231 cells treated with nocodazole (Supplementary Fig. 2a; Supplementary Datasets 6-9).

**Figure 2.**
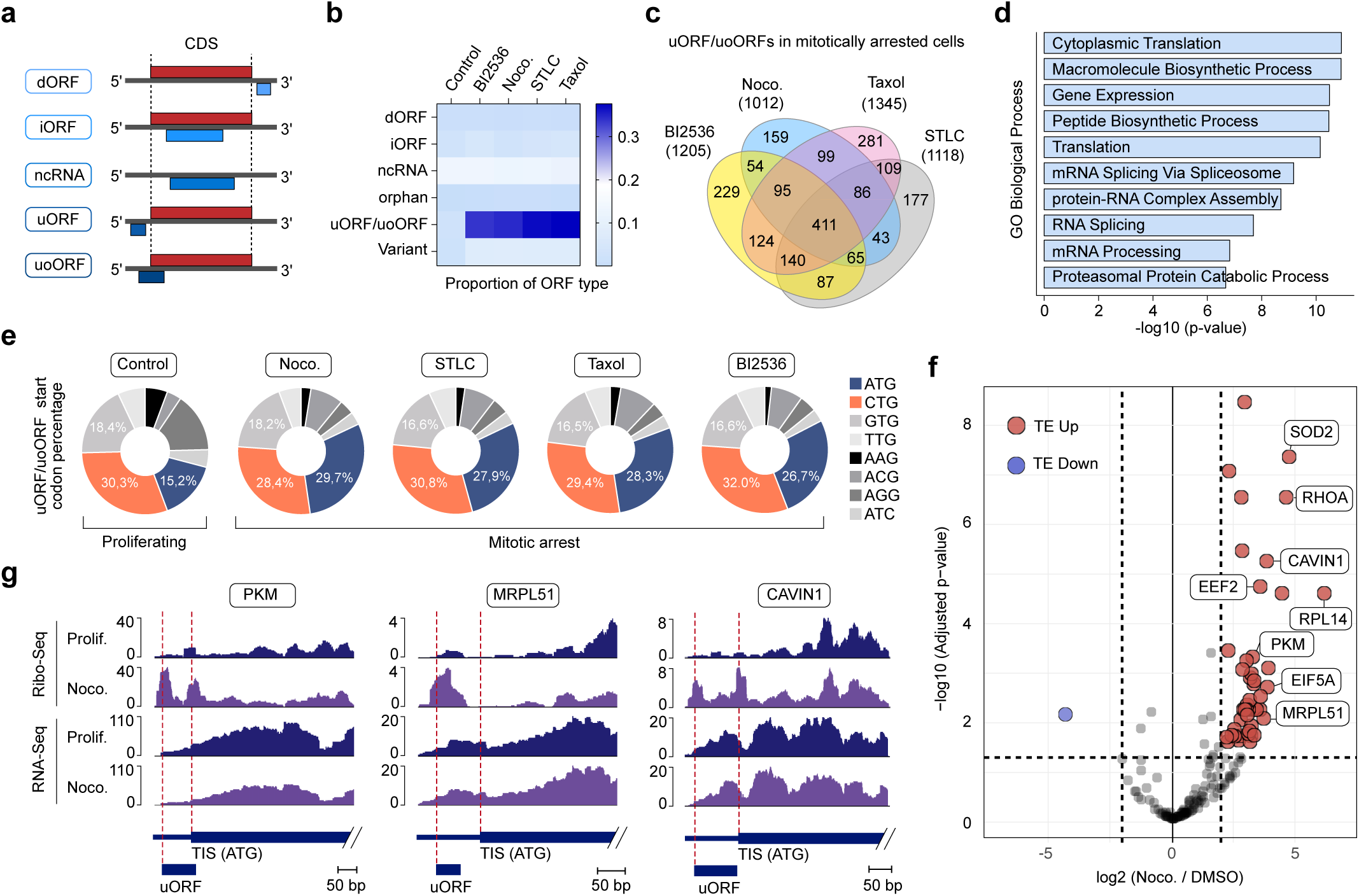
Increased translation of uORFs/uoORFs in mitotically arrested cancer cells. **a**, Schematic overview of ORF types detected by PRICE. dORF, downstream open reading frame; iORF, internal open reading frame; ncRNA, non-coding RNA. **b,** Proportion of ORFs of each type identified in U2OS cells treated with vehicle (Control), BI2536 (0.1 µM), Nocodazole (0.5 µM), Taxol (1 µM), or STLC (1 µM) for 16 hours. "Variant" refers to ORFs that contain a stop codon but cannot be categorized as CDS or truncated CDS. "Orphan" refers to ORFs that do not fit into any of the described categories. **c,** Venn diagram showing the number of uORFs/uoORFs identified by PRICE in U2OS cells arrested in mitosis with BI2536 (0.1 µM), Nocodazole (0.5 µM), Taxol (1 µM), or STLC (1 µM) for 16 hours. **d,** Bar plot showing the combined scores for the GO database "Biological Process" obtained by Enrichr analysis of the common genes described in panel c. **e,** Percentage of translation initiation site codons from uORFs/uoORFs in U2OS cells treated with vehicle (Control), BI2536 (0.1 µM), Nocodazole (0.5 µM), Taxol (1 µM), or STLC (1 µM) for 16 hours. **f,** Volcano plot illustrating the translation efficiency (TE) of all predicted uORFs/uoORFs in U2OS cells treated with 0.5 µM Nocodazole for 16 hours compared to asynchronous cells. The x-axis represents the log2 fold change in TE for uORFs/uoORFs. The y-axis indicates the significance of changes in TE. uORFs/uoORFs with increased TE are highlighted in red, while those with decreased TE are highlighted in blue. A chi-square test resulted in a p-value of 5.29e-42, indicating a significant overrepresentation of uORF/uoORFs within genes exhibiting differential TE. **g,** Read distribution of representative uORFs/uoORFs with increased TE in mitotically arrested U2OS cells. Ribo-Seq (upper panels) and RNA-Seq (lower panels) reads are shown for the 5’UTR and the start of the CDS. Prolif., proliferating; Noco., Nocodazole.

More than 80% of the over 1,000 uORFs/uoORFs identified in mitotically arrested U2OS cells were commonly induced by at least two distinct molecular mechanisms (Fig. 2c). The uORFs/uoORFs exclusively translated in mitotically arrested cells were enriched in genes involved in biological processes such as cytoplasmic translation, mRNA splicing, and proteasome-mediated catabolic processes (Fig. 2d). These genes were also associated with cellular components like the cytoskeleton and focal adhesions (Supplementary Fig. 2b). Interestingly, the majority of predicted uORFs/uoORFs in proliferating U2OS cells initiated from non-ATG start codons (∼84%). In contrast, a decrease to ∼72% was observed in mitotically arrested cells, indicating a shift toward more canonical ATG initiation sites. Notably, the total number of uORFs/uoORFs with ATG initiation sites increased by approximately 80% in mitotically arrested cancer cells (Fig. 2e; Supplementary Fig. 2c).

To assess whether translation rates of these predicted uORF/uoORFs were elevated in mitotically arrested cells, we calculated their translation efficiencies (TE) by normalizing RiboSeq reads mapped to genomic uORF/uoORFs coordinates against RNASeq reads from the same regions. Our analysis revealed that the vast majority of predicted uORF/uoORFs exhibited increased TE in mitotic cells (Fig. 2f). Notable examples included PKM, MRPL51, and CAVIN1, which are implicated in critical cellular processes such as energy metabolism, mitochondrial function, and the oxidative stress response (Fig. 2g). Furthermore, mapping translation initiation sites (TIS) using Harringtonine in mitotically arrested cells demonstrated clear ribosome footprint peaks at predicted uORF/uoORFs start sites (Supplementary Fig. 2d). Collectively, our data indicate that ribosome redistribution during mitosis significantly enhances the translation of hundreds of uORF/uoORFs.

### Immunopeptidomics identifies uORF/uoORF-derived peptides presented by HLA Class I in cancer cells

Next, we investigated the presentation of uORF/uoORF-derived peptides on the surface of cancer cells via human leukocyte antigen (HLA) molecules. We employed state-of-the-art liquid chromatography-tandem mass spectrometry (LC-MS/MS)-based immunopeptidomics on U2OS cells and the triple-negative breast cancer (TNBC) cell line SUM-159PT. For each cell line, three biological replicates of 5×10⁷ cells were arrested in mitosis using Taxol, a common first-line therapy in metastatic breast cancer. This approach allowed for the sensitive, high-throughput identification of HLA-I-presented peptides. We first constructed a comprehensive mitotic uORF/uoORFs database (uORF/uoORFdb) by selecting and translating nucleotide sequences of all predicted uORF/uoORFs in the cell lines used in this study. This database, along with the annotated proteome, served as a reference for our MS-immunopeptidomics data analysis (Fig. 3a)

**Figure 3.**
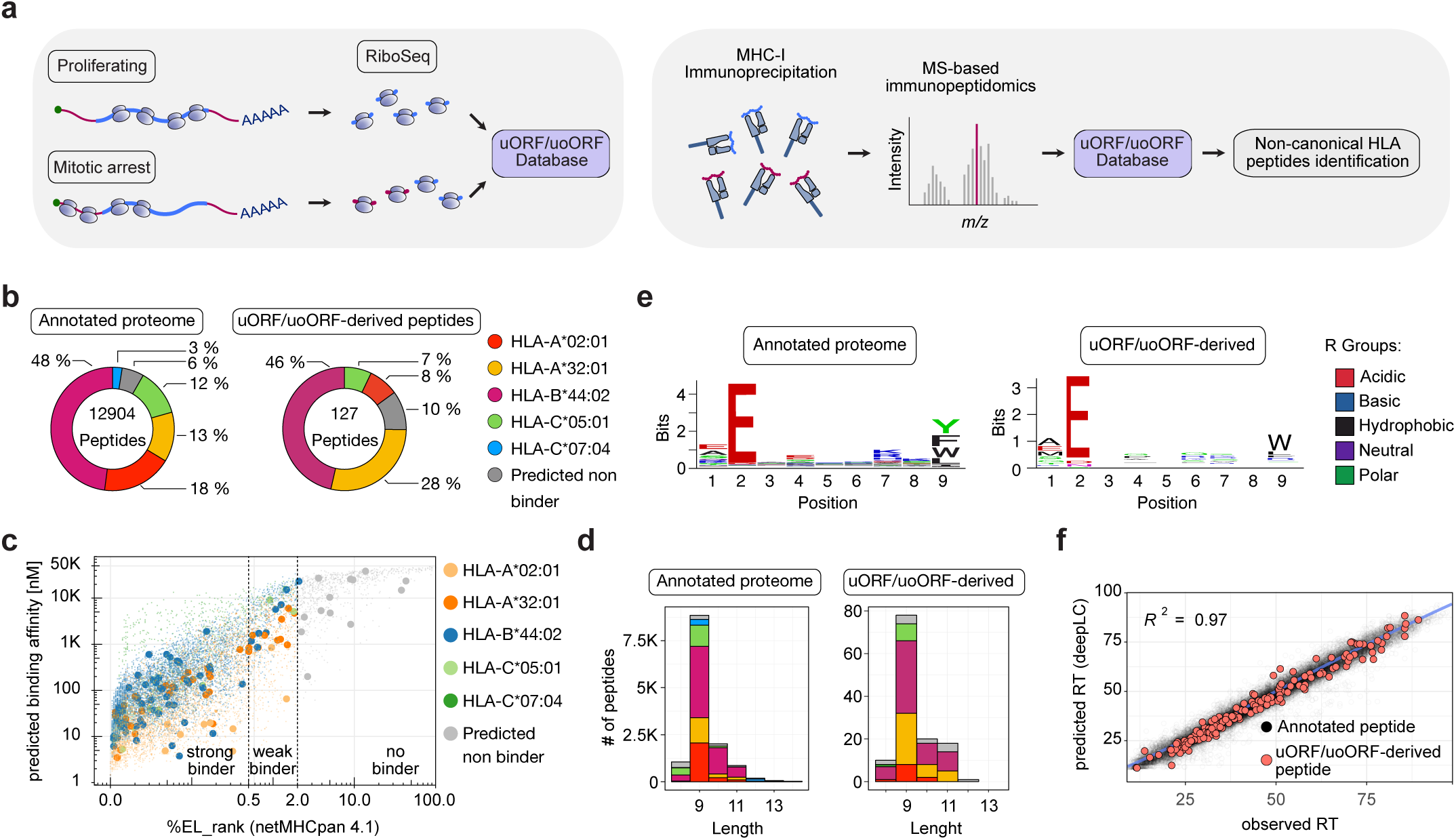
uORF/uoORF-derived HLA-presented peptides exhibit comparable characteristics to annotated peptides. **a**, Schematic overview of the uORF/uoORF database creation using Ribo-seq and PRICE prediction, followed by the identification of non-canonical HLA peptides through liquid chromatography-tandem mass spectrometry (LC-MS/MS)-based immunopeptidomics. **b,** Peptide quantification in the annotated proteome (left panel) and uORF/uoORF-derived proteome (right panel) of proliferating and mitotically arrested U2OS cells. The distribution of predicted binding to HLA alleles in U2OS cells is shown. **c,** Percentage of eluted ligand (EL) peptides predicted by NetMHCpan-4.1 plotted against predicted binding affinity for peptides derived from the annotated proteome (small dots) and uORF/uoORF-derived peptides (large dots) in proliferating and mitotically arrested U2OS cells. Predicted binding to HLA alleles in U2OS cells is shown. Peptides are categorized as strong binders (%EL rank 0–0.5), weak binders (%EL rank 0.5–2), or non-binders (%EL rank 2–100). **d,** Length distribution of detected peptides for the annotated proteome (left panel) and uORF/uoORF-derived peptides (right panel) in proliferating and mitotically arrested U2OS cells. The proportion of predicted binding to U2OS HLA alleles is shown. **e,** Peptide motif plots of unique peptides from the annotated proteome (6,624) and unique peptides derived from uORFs/uoORFs (58), confidently identified as binding to the U2OS allele HLA-B44:02 **f,** Observed retention time (RT) indices plotted against predicted RT indices for peptides from the annotated proteome (black) and uORF/uoORF-derived proteome (red) across all HLA alleles in proliferating and mitotically arrested U2OS cells. R² represents the Pearson correlation coefficient.

Our uORF/uoORFdb comprised 9,008 predicted ORFs (Supplementary Dataset 10), representing a significant expansion from previous studies. This extensive database allowed us to screen for more than 6,700 uORFs and over 2,200 uoORFs, substantially increasing the likelihood of identifying novel non-canonical ORF-derived HLA-presented peptides. Notably, our approach aligns with recent findings that uORF/uoORFs can encode biologically active proteins and HLA-presented peptides in both malignant and benign cells, suggesting their potential role in cancer cell development and survival^35^.

Our immunopeptidomics analysis revealed a substantial repertoire of HLA class I-presented peptides in U2OS and SUM-159PT cells, with 12,904 and 25,655 unique proteome-derived peptides identified, respectively. Notably, we discovered 127 and 166 uORF-derived HLA class I-presented peptides in these cell lines (Fig. 3b; Supplementary Fig. 3a), representing 0.5–1% of the MHC-I immunopeptidome in mitotically arrested cells.

Binding prediction revealed similar distribution of assigned alleles for peptides derived from the canonical proteome and uORF/uoORF-derived peptides (Fig. 3b; Supplementary Fig. 3a). Predicted binding affinities to different HLA allotypes showed that 91% of uORF/uoORF- derived peptides and 90% of proteome-derived peptides were likely to bind to the HLA allotypes (Fig. 3c; Supplementary Fig. 3b). Both types of peptides, those from the canonical proteome and predicted uORF/uoORFs, exhibited the typical length distribution of HLA class I-presented peptides, predominantly as nonamers (Fig. 3d; Supplementary Fig. 3c). Sequence clustering of these peptides allowed reconstruction of the binding motifs for the HLA allotypes expressed by the respective cell line (Fig. 3e; Supplementary Fig. 3d). Moreover, predicted retention times for uORF/uoORF-derived peptides showed a high correlation with observed chromatographic retention times, comparable to those of annotated peptides (Fig. 3f; Supplementary Fig. 3e). Overall, the quality of uORF/uoORF-derived peptide identifications was comparable to that of annotated peptides, supporting their authenticity as genuine HLA class I-presented peptides. In summary, our results provide evidence that uORF/uoORF- derived peptides are presented by HLA class I molecules, suggesting that they may represent an underappreciated source of tumor antigens and potentially expand the repertoire of targets for cancer immunotherapy. This finding aligns with recent studies that underscore the significance of non-canonical peptides in the immunopeptidome^35^.

### Identification of therapy-induced uORF/uoORF-derived peptides as potential antigenic targets in mitotically arrested cancer cells

Mitotic inhibitors offer a promising option for combination with checkpoint inhibitors or other immunotherapies to enhance immune response effectiveness^36^. However, the impact of mitotic arrest on the HLA class I peptide repertoire remains uncharacterized. To address this, we aimed to quantify changes in uORF/uoORF-derived peptides in cancer cells following mitotic arrest *in vitro*, to better understand how mitotic inhibition could be leveraged in combination therapy regimens to improve patient outcomes.

To assess MHC-I repertoire alterations induced by mitotic arrest, we performed label-free quantification of HLA-presented peptides, comparing Taxol-treated cells to DMSO-treated controls in U2OS and SUM-159PT cell lines. We identified 13 uORF/uoORF-derived peptides enriched in the immunopeptidome of mitotic U2OS cells and 25 candidates in mitotically arrested SUM-159PT cells (Fig. 4a-b; Supplementary Fig. 4a-b). Notably, these candidates showed increased translation of their respective uORFs/uoORFs, while RNA levels remained unchanged (Fig. 4c). Additionally, mapping translation initiation sites with harringtonine revealed an enrichment of ribosome-protected fragments at upstream initiation sites in mitotic cells (Supplementary Fig. 4c). Importantly, none of the identified peptide sequences were found in the HLA Ligand Atlas.

**Figure 4.**
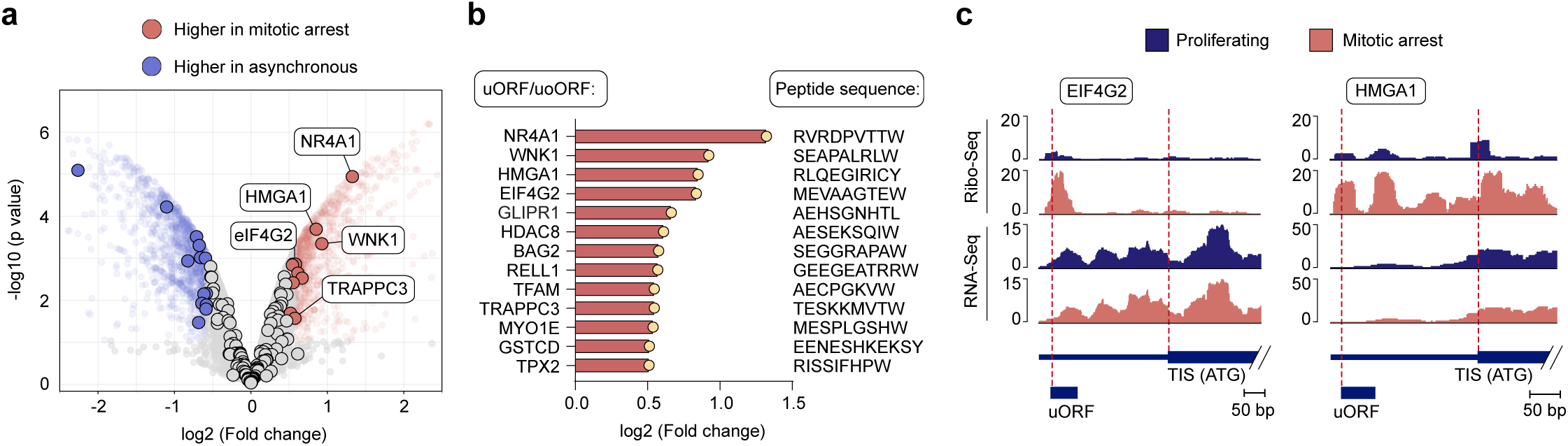
Quantitative analysis of HLA-presented uORF/uoORF-derived peptides in mitotically arrested U2OS cells. **a**, Volcano plot illustrating the label-free quantification of the immunopeptidome in Taxol-treated versus DMSO-treated U2OS cells, with genes expressing uORF/uoORF-derived peptides highlighted. Peptides with a log2 fold change > 0.5 and adjusted p-value < 0.05 are marked in red, while those with a log2 fold change < -0.5 and adjusted p-value < 0.05 are marked in blue. Statistical analysis was performed using an empirical Bayes moderated *t*-test with two-sided *p*-values. **b,** Name of the gene expressing the uORF/uoORF-derived peptide, log2 fold change (FC) of peptide abundance, and peptide sequence of mitotic arrest-induced peptides in U2OS cells. **c,** Read distribution of representative uORFs/uoORFs with increased expression in mitotically arrested U2OS cells. Ribo-Seq (upper panels) and RNA-Seq (lower panels) reads are displayed for the 5’ UTR and the start of the coding sequence (CDS).

The genes expressing the upregulated uORF/uoORF immunopeptidome in mitotically arrested U2OS cells were associated with cellular components like the cytoskeleton and adherens junctions (Supplementary Fig.4d). One of the detected peptides stems from a uORF in the *eIF4G2* gene, a non-canonical translation initiation factor that plays a crucial role in mitosis by facilitating the translation of specific mRNAs essential for cell division, including CDK1, which regulates phase transitions in the cell cycle^37^. Additionally, we identified a uORF-derived peptide from the *HMGA1* gene, a protein that significantly influences mitosis by modulating gene expression and chromatin structure. During cell division, HMGA1 promotes the transcription of genes involved in cell cycle progression, particularly those associated with the G2/M transition^38,39^. These newly characterized, uORF-derived peptides, being clearly enriched in mitotically arrested cells, represent promising antigenic targets for immunotherapy, aligning with recent findings that emphasize the role of non-canonical peptides in shaping immune responses.

### Enhanced presentation of uORF-derived peptides in mitotically arrested cancer cells facilitates targeted immune responses

The therapy-induced expression of uORF/uoORF-derived peptides on the surface of cancer cells presents a promising opportunity for targeted immunotherapy. These peptides, displayed by cancer cells arrested in mitosis following chemotherapy, may offer a unique immunogenic signature, making them potential candidates for precise immune targeting. To explore this, we first generated luciferase reporters containing 5’ UTRs with uORFs that showed increased peptide presentation in mitotically arrested cells. The peptide sequences identified by immunopeptidomics were replaced with the SIINFEKL peptide from chicken ovalbumin (OVA), an 8-amino acid peptide presented by H2-K^b^. This complex is recognized by T cell receptor (TCR)-transgenic CD8^+^ OT-I T cells, as well as by a TCR-like antibody (clone 25-D1.16), which allows for assessment of MHC presentation of the SIINFEKL epitope via flow cytometry^40^. For this analysis, we specifically selected the 5’ UTRs of eIF4G2 and TPX2 (Fig. 5a).

**Figure 5.**
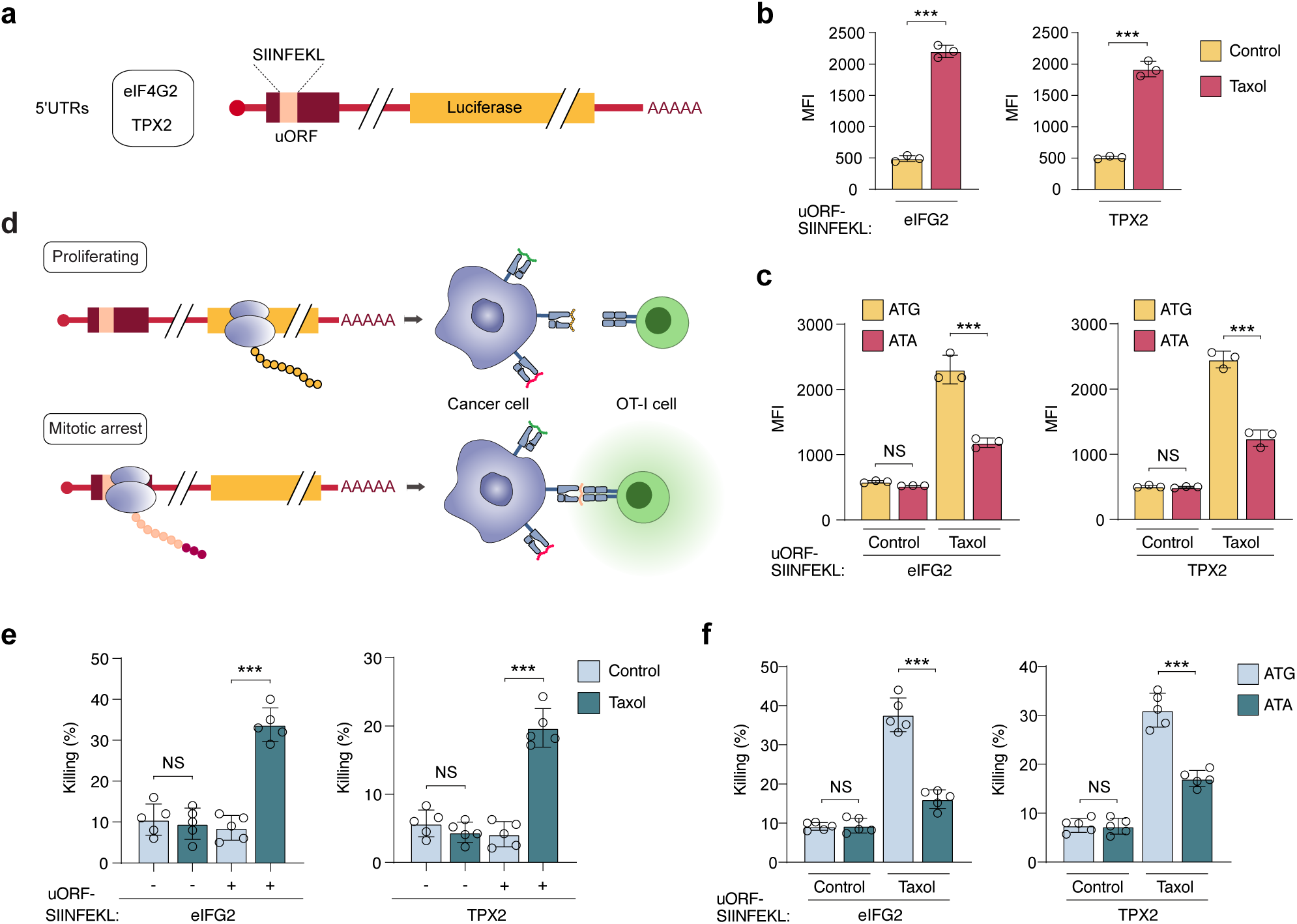
Increased uORF-derived peptide presentation in mitotic cancer cells promotes targeted immune responses. **a**, Schematic of the uORF reporter system. Two different 5’ UTRs (eIF4G2 and TPX2) were selected and cloned into the pGL3-promoter vector. The sequence of the uORF-derived peptide was replaced with the SIINFEKL sequence, followed by firefly luciferase. **b,** Detection of the p:MHC complex SIINFEKL:H-2K^b^ by flow cytometry, shown as Median fluorescence intensity (MFI), in TC1 cells transfected with the indicated reporters. Cells were treated with Taxol (1 µM) or DMSO (Control) for 16 hours. Data represent mean ± SD from biologically independent experiments (*n =* 3). *p*-values were calculated using a two-tailed unpaired *t*-test. \*\*\**P* < 0.001. **c,** MFI of the SIINFEKL peptide bound to H-2K^b^ in TC1 cells transfected with wild-type uORF-SIINFEKL reporters or start codon mutant uORF-SIINFEKL reporters. Cells were treated with Taxol (1 µM) for 16 hours. Data represent mean ± SD from biologically independent experiments (*n =* 3). *p*-values were calculated using a two-tailed unpaired *t*-test. \*\*\**P* < 0.001. **d,** Schematic illustrating how the presentation of uORF-derived SIINFEKL peptides induced by mitotic arrest leads to OT-I T cell recognition. Proliferating cells expressing uORF-SIINFEKL reporters are not recognized by OT-I cells, whereas enhanced uORF translation in mitotically arrested cells allows for their recognition. **e-f**, Percentage of TC1 cell killing *in vitro* by activated OT-I cells. TC1 cells were transfected with the indicated reporters and arrested in mitosis with Taxol (1 µM) for 16 hours or treated with DMSO for the same period. Data represent mean ± SD from biologically independent experiments (*n =* 5). *p*-values were calculated using a two-tailed unpaired *t*-test. NS, non-significant; \*\*\**P* < 0.001.

Next, we evaluated whether therapy-induced mitotic arrest enhances the presentation of uORF-derived peptides in cancer cells. Transfection of uORF-SIINFEKL reporters into the murine cancer cell line TC-1 revealed that reporter expression in proliferating cells did not result in significant recognition by the 25-D1.16 antibody. In contrast, mitotically arrested cells exhibited a marked increase in uORF-derived SIINFEKL presentation, regardless of the agent used to induce the arrest (Fig. 5b; Supplementary Fig.5a). Importantly, mRNA levels of the reporters were not affected during mitotic arrest (Supplementary Fig. 5b). To further investigate the role of uORF translation during mitotic arrest, we mutated the uORF start codons from ATG to ATA. TC-1 cells expressing these mutant reporters showed reduced SIINFEKL presentation upon mitotic arrest, indicating that active uORF translation during mitosis is crucial for effective antigen presentation (Fig. 5c).

To test whether the increased presentation of uORF-derived peptides during mitotic arrest enhances CD8^+^ T cell-mediated cytotoxicity, we activated OT-I CD8^+^ T cells with anti-CD3, anti-CD28, and IL-12 for 72 hours and co-cultured them with either proliferating or mitotically arrested TC-1 cells expressing the uORF-SIINFEKL reporters (Fig. 5d). Neither transfection nor mitotic arrest alone resulted in significant cytotoxic effects. However, mitotic arrest in reporter-expressing cells significantly enhanced cytotoxic activity and tumor cell killing by OT-I T cells *in vitro* (Fig. 5e). Consistent with these observations, OT-I T cells secreted IFN-γ only after co-culture with mitotically arrested cancer cells expressing the uORF-SIINFEKL reporters (Supplementary Fig. 5c). Notably, cancer cells expressing reporters with mutant start codons showed reduced antigen-specific killing by OT-I T cells, accompanied by a significant decrease in IFN-γ secretion during co-culture (Fig. 5f; Supplementary Fig.5d). These findings highlight the potential of therapy-induced uORF-derived peptides as innovative targets for immunotherapy, underscoring their role in enhancing immune recognition during cancer treatment.

## Discussion

In this study, we demonstrate that cancer cells treated with mitotic inhibitors, such as Taxol or Nocodazole, undergo a significant redistribution of ribosomes toward the 5’UTR and the start of the coding sequence. Cells arrested in mitosis exhibit elevated translation initiation, driven by the sustained phosphorylation of 4E-BP1 by CDK1^40,41^. This results in increased initiation at non-canonical sites within the 5’ UTR. By applying bioinformatic prediction tools, we identified that these non-canonical initiation events lead to the translation of thousands of novel peptides originating from upstream open reading frames (uORFs/uoORFs). Additionally, through immunopeptidomics, we showed that dozens of these non-canonical peptides are presented by HLA class I molecules on the surface of cancer cells. Notably, the presentation of a model therapy-induced neoepitope allowed T lymphocytes to specifically recognize and kill drug-treated cancer cells. Importantly, none of the uORF/uoORF-derived neoepitopes were found in existing human immunopeptidomics or proteomics databases, underscoring their treatment-specific nature. We have termed these newly identified sequences as "therapy-induced uORF/uoORF-derived epitopes."

These therapy-induced peptides present a promising opportunity for the development of personalized vaccines or therapies that elicit robust immune responses specifically targeting chemotherapy-treated cancer cells. Paclitaxel (Taxol), in particular, emerges as a strong candidate for combinatorial T cell-based immunotherapies. Beyond its well-known role as a microtubule stabilizer, and its new role as a uORF/uoORF-derived epitope inducer described in this work, paclitaxel exhibits significant immunomodulatory effects that enhance tumor immunogenicity. Notably, it induces immunogenic cell death (ICD), which leads to the release of tumor antigens and damage-associated molecular patterns (DAMPs)^41,42^. This process bolsters the antigen-presenting capabilities of dendritic cells (DCs) and macrophages, thereby amplifying anti-tumor immune responses. Additionally, paclitaxel promotes Th1 cellular immunity by increasing the levels of IFN-γ-secreting CD8^+^ T cells and IL-2-secreting CD4^+^ T cells^43^. This dual activation enhances both cytotoxic and helper T cell responses, crucial for effective tumor clearance. Moreover, paclitaxel has been shown to inhibit immunosuppressive cells, such as myeloid-derived suppressor cells (MDSCs) and regulatory T cells (Tregs), further potentiating the overall anti-tumor immune response^43^.

During mitosis, mammalian cells increase the stringency of start-codon selection, largely through the release of nuclear eIF1, which alters the translation of numerous proteins and helps preserve cell viability under mitotic stress, especially during delays induced by anti- mitotic drugs^44^. These altered translation events potentially lead to synthesis of thousands of non-canonical peptides. Historically, non-canonical open reading frames (ncORFs) have been excluded from genome annotations due to concerns about false positives and the lack of robust methods for validating their translation^45^. However, recent advances in ribosome profiling (Ribo-Seq) have revolutionized our understanding of ncORFs by enabling the detection of actively translated regions at near-codon resolution. This has led to a more comprehensive view of the translatome. Several bioinformatics tools, including PRICE^34^, RibORF^46^, ORFquant^47^, or Ribotricer^48^, analyze Ribo-Seq data to identify ncORFs based on the characteristic 3-nucleotide periodicity of translating ribosomes. These tools have revealed that thousands of ncORFs are translated in human cells, suggesting they play a previously underappreciated role in expanding both the human proteome and immunopeptidome^49^.

In particular, ncORF-derived peptides induced by therapeutic interventions represent a largely untapped source within the HLA class I immunopeptidome. Identifying and characterizing these therapy-induced peptides could offer novel, therapy-specific targets for immunotherapy, potentially opening new avenues for personalized treatments^50,51^. Notably, uORF/uoORF- derived peptides may even be shared across different tumor types and between patients, suggesting their potential as common targets for broader, cross-entity immunotherapeutic approaches. This shared presence of uORF/uoORF-derived peptides could enhance the relevance of these non-canonical peptides in developing universal or semi-personalized cancer immunotherapies, further expanding the impact of immunopeptidome research.

## Methods

### Cell culture

All cell lines were cultured in DMEM High Glucose (ThermoFisher Scientific) supplemented with 10 % FBS (ThermoFisher Scientific) and 1 % PenStrep (ThermoFisher Scientific) at 37°C and 5 % CO2. For mitotic shake-off experiments, cells were cultured to a confluency of 70% and treated with mitosis-arresting compounds for 16 hours; BI2536, 0.1 µM (Cell Signaling Technology, #26744), Nocodazole, 0.5 µM (Sigma-Aldrich, M1404), Taxol, 1 µM (Santa Cruz, sc-201439), STLC 5 µM (Tocris, #2191). The medium was centrifuged 600x G for 10 min. at 4 °C. The cell pellets were washed in ice-cold PBS. All cell lines were regularly tested for Mycoplasma contamination.

### Propidium iodide (PI) staining

Cells were harvested as previously described, resuspended in PBS containing 1 mM EDTA, and counted using a Casy Counter system. The cell concentration was adjusted to ensure equal cell numbers. The cells were then fixed in ice-cold absolute ethanol and stored overnight at -20°C. Following storage, the cells were washed three times with PBS + 1 mM EDTA and resuspended in PI staining buffer (50 µg/ml PI, 0.2 mg/ml RNase A, 0.4% Triton X-100, 1 mM EDTA in PBS). The cells were incubated at 37°C for 30 minutes with constant shaking. After incubation, the samples were washed with PBS + 1 mM EDTA, filtered into round-bottom tubes containing cell strainers, and analyzed using a BD Canto II system. To select PI-stained single cells, the following gating strategy was applied: cells were initially separated using FSC and SSC to distinguish the cell population. Single cells were then identified by combining FSC-H and FSC-A, followed by 488 nm excitation and filtering for the BL84/42 signal.

### Immunoblotting

Cell pellets were harvested and lysed in whole-cell lysis buffer (1M Tris pH 7.5, 10 % glycerol, 2 % SDS). Next, 1 µl of 0.1 M MgCl2 solution and 2.5 U of Benzonase (Merck) was added and incubated at 37 °C for 10 min to digest DNA. Bradford assay (Serva, Bradford reagent 5x) was used to determine and level protein input. Lysates were diluted with water and 4x Laemmli Buffer (BioRad) supplemented with 10 % β-mercaptoethanol. Samples were separated by SDS-PAGE and transferred to nitrocellulose membranes using TurboBlot (BioRad) with transfer-buffer (240mM Tris-HCl, 195mM Glycin, 0.5% SDS). Next, membranes were blocked in TBS-T plus either 5% non-fat dry milk or 5% BSA for 1 hour at room temperature. Primary antibodies were diluted in TBS-T plus either 5% non-fat dry milk or 5% BSA and incubated with the membrane overnight at 4°C and gentle rotation. Next, membranes were washed 4 times with TBS-T for 5 min at room temperature before incubating with the secondary antibody in TBS-T for 1 hour at room temperature, protected from light. Membranes were imaged using an Odyssey CLx machine (LICORbio). Antibodies used were 4E-BP1 (Cell Signaling; cat. #9644, 1:1,000), Phospho-4E-BP1 (Thr37/46) (Cell Signaling; cat. #2855, 1:1,000), Phospho-Histone H3 (Ser10) (Cell Signaling; cat. #9701, 1:1,000), GAPDH (ProteinTech; cat. #60004; 1:20,000).

### RiboSeq

Cells were resuspended in lysis buffer (20 mM Tris-HCl, pH 7.5, 10 mM MgCl₂, 100 mM KCl, 1% Triton X-100), supplemented with 100 µg/mL cycloheximide (CHX), 2 mM DTT, and 1x Complete Protease Inhibitor (Roche). They were then treated with RNase I (Ambion) at a concentration of 1.2 U/µL for 45 minutes at room temperature. The samples were layered onto sucrose gradients, ranging from 47% to 7% sucrose, prepared in 20 mM Tris-HCl, pH 7.5, 10 mM MgCl₂, 100 mM KCl, 100 µg/mL CHX, and 2 mM DTT. The gradients were centrifuged for 2 hours at 36,000 rpm at 4 °C using a Beckman-Coulter ultracentrifuge with an SW41-Ti rotor. Monosome-containing fractions were collected and digested with 15 µL of recombinant Proteinase K (Roche) and 1% SDS for 45 minutes at 45 °C. Subsequently, RNA was extracted using a standard phenol-chloroform-guanidinium thiocyanate method. The resulting ribosome-protected fragments (RPFs) were size-selected (20-34 nt) and 3’ dephosphorylated using T4 Polynucleotide Kinase (PNK; New England Biolabs). Following this, 5’ pre-adenylated linkers were ligated to the 3’ ends of the RPFs using T4 RNA Ligase 2 (New England Biolabs), and rRNA depletion was performed using custom biotinylated rRNA-oligonucleotides and streptavidin-coated magnetic beads. Reverse transcription was carried out using the SuperScript III First-Strand Synthesis Kit (Thermo Fisher). cDNA was circularized using CircLigase ssDNA Ligase II (LGC Biosearch Technologies) and then amplified via PCR using Q5 High-Fidelity 2x Master Mix (New England Biolabs). DNA was quantified using the Qubit dsDNA HS Kit and adjusted to a concentration of 2 nM, suitable for NextSeq 2000 sequencing. All oligo sequences used in this study are listed in Supplementary Dataset 7.

### Harringtonine assay

Cells were treated with 2 µg/mL Harringtonine at the indicated time points before harvest. Control cells were scraped into the medium, while mitotically arrested cells were harvested by shake-offs and collected from the medium. All samples were pelleted and resuspended in lysis buffer (20 mM Tris-HCl, pH 7.5, 10 mM MgCl₂, 100 mM KCl, 1% Triton X-100), supplemented with 100 µg/mL cycloheximide (CHX), 2 mM DTT, and 1x Complete Protease Inhibitor (Roche). Thus, all samples were exposed to the same duration of Harringtonine treatment.

### RiboSeq Analysis

Sample adapters were trimmed using cutadapt (v3.4) and demultiplexed with barcode_splitter from FASTX-toolkit (v0.0.6). Fragments smaller than 20 nt were dropped. UMIs extraction was performed using umi_tools (v1.1.1). By BLAST-Like Alignment Tool (BLAT) (v36x2), rRNA reads were filtered and discarded. The rRNA index for RNA18S5, RNA28S5 and RNA5-8S5 was constructed manually from NCBI RefSeq annotation. Remaining reads were aligned with Spliced Transcripts Alignment to a Reference (STAR) (v2.5.3a) to GRCh37/hg19 with -- outSAMtype BAM Unsorted --readFilesCommand zcat --quantMode TranscriptomeSAM GeneCounts --outSAMmapqUnique 0. Genome browser bigwig tracks were obtained using samtools (v1.15.1) and bedtools (v2.24.0).

### Transcript distribution

To analyze ribosome transcript distribution, we selected the most representative isoform for each gene using a hierarchical selection strategy. RPF counts were obtained for each selected transcript, and intra-gene normalization was performed by dividing the cumulative read counts for each region (5’ UTR, CDS, 3’ UTR) by the total RPF counts for that transcript. This normalization allowed for comparisons across different regions of the same transcript. Next, we computed RPF density across regions for each transcript by interpolating read counts over a fixed grid of 2,000 points. Transcripts with fewer than 50 reads were excluded from the downstream analysis. The interpolated RPF densities across transcripts were then averaged and subjected to Gaussian smoothing to reduce noise. Finally, the resulting RPF densities were plotted alongside the corresponding transcript regions. All analyses were performed using custom Python scripts that incorporated the NumPy, SciPy, and Matplotlib libraries.

### Upstream translation initation sites quantification

Predicted uORF and uoORF genomic coordinates from U2OS RiboSeq data were compiled in SAF format and counted using featureCounts from the subread package (v1.5.1). Resulting counting tables were filtered for >= 5 reads per sample and feature. Next, trimmed mean of M-values (TMM) for cross-sample comparison were determined using the "calcNormFactors" function from edgeR. Normalized counts per million (cpm) were subjected to sample-specific outlier calculation using Grupps function Log2(TMM) values were calculated using a custom awk script.

### ORF Prediction

ORFs were predicted using PRICE^50^. In brief, UMI-extracted and rRNA-filtered FASTQ files were re-aligned to the GRCh37/hg19 reference genome using STAR (v2.5.3a). Important outSAMattributes required by PRICE were specified, including --outSAMtype BAM Unsorted, --alignEndsType Extend5pOfReads12, --outSAMattributes nM MD NH, --readFilesCommand zcat, --quantMode TranscriptomeSAM GeneCounts, and --outSAMmapqUnique 0. The PRICE reference genome was prepared as described, utilizing hg19 FASTA and GTF files from Gencode. Next, PRICE was executed with the respective BAM files using the command ∼/Gedi/Gedi_1.0.5/gedi -e Price -D -genomic hg19 -progress -plot. Subsequently, all ORF features with a p-value of p ≤ 0.05 were quantified using standard UNIX commands. ORF tables from PRICE were adjusted to the BED format, including chromosome, ORF feature start position, ORF feature end position, ORF feature ID, chromosome strand, and Gene ID. The resulting BED6 files were converted to BED12 format, which served as input for bedtools getfasta with the flags -s, -name, and -split. Peptide sequences were generated using the faTrans program.

### Translational efficiency

Translational efficiency of uORF/uoORFs was calculated by extracting IDs, start and end positions of predicted uORF and uoORF features (pvalue < 0.05) from PRICE ORF tables and arranged in SAF format, creating a uORF/uoORF SAF reference file. Next, read counts in uORF/uoORF regions were determined with featureCounts (v1.5.1) and genome-based BAM files from RNAseq and RiboSeq. The resulting aggregated count matrix was subjected to RiboDiff (v0.2.1) calculation. Features with missing calculation were discarded. Data was generated using pseudoreplicates to ensure robustness in translational efficiency estimates.

### Immunopeptidomics

Input cell lines (U2OS and SUM159) for immunopeptidomics were harvested in triplicates with 5×10^7^ cells per replicate. The cell lines U2OS and SUM-159PT were treated with 1 µM Taxol for 16 hours. For DMSO-treated control conditions, cells were gently scraped in ice-cold PBS. For Taxol-treated conditions, mitotic shake-offs were spun at 600x g for 10 min at 4 °C and washed with ice-cold PBS. All samples of all conditions were counted using a CASY II system and snap-frozen.

Immunoprecipitation of HLA class I:peptide complexes was performed as previously described^52^ with additional steps for the forced oxidation of methionine using H_2_O_2_ and reduction and alkylation of cysteine using tris(2-carboxyethyl)phosphine (TCEP) and iodoacetamide (IAA).

Lyophilized peptides were dissolved in 12 µl of 5% ACN in 0.1% TFA and spiked with 0.5µl of 100 fmol/µl Peptide Retention Time Calibration (PRTC) Mixture (Pierce) and 10 fmol/µl JPTRT 11 (a subset of peptides from the Retention Time Standardization Kit; JPT) and transferred to QuanRecovery Vials with MaxPeak HPS (Waters, Milford, MA, USA). All samples were analyzed using an UltiMate 3000 RSLCnano system coupled to an Orbitrap Exploris 480 equipped with a FAIMS Pro Interface (Thermo Fisher Scientific). For chromatographic separation, peptides were first loaded onto a trapping cartridge (Acclaim PepMap 100 C18 μ- Precolumn, 5μm, 300 μm i.d. x 5 mm, 100 Å; Thermo Fisher Scientific) and then eluted and separated using a nanoEase M/Z Peptide BEH C18 130A 1.7µm, 75µm x 200mm (Waters). Total analysis time was 120 min and separation was performed using a flow rate of 0.3 µl/min with a gradient starting from 1% solvent B (100% ACN, 0.1% TFA) and 99% solvent A (0.1% FA in H_2_O) for 0.5 min. Concentration of solvent B was increased to 2.5% in 12.5 min, to 28.6% in 87 min and then to 38.7% in 1.4 min. Subsequently, concentration of solvent B was increased to 80% in 2.6 min and kept at 80% solvent B for 5 min for washing. Finally, the column was re-equilibrated at 1% solvent B for 11 min. The LC system was coupled on-line to the mass spectrometer using a Nanospray-Flex ion source (Thermo Fisher Scientific), a SimpleLink Uno liquid junction (FossilIonTech) and a CoAnn ESI Emitter (Fused Silica 20 µm ID, 365 µm OD with orifice ID 10 µm; CoAnn Technologies). The mass spectrometer was operated in positive mode and a spray voltage of 2400 V was applied for ionization with an ion transfer tube temperature of 275 °C. For ion mobility separation, the FAIMS module was operated with standard resolution and a total carrier gas flow of 4.0 l/min. Each sample was injected twice using either a compensation voltage of -50 V or -65 V for maximal orthogonality and thus increased immunopeptidome coverage. Full Scan MS spectra were acquired for a range of 300 – 1650 m/z with a resolution of 120.000 (RF Lens 50%, AGC Target 300%). MS/MS spectra were acquired in data-independent mode using 44 previously determined dynamic mass windows optimized for HLA class I peptides with an overlap of 0.5 m/z. HCD collision energy was set to 28% and MS/MS spectra were recorded with a resolution of 30.000 (normalized AGC target 3000%).

MS raw data was analyzed using Spectronaut software (version 17.6; Biognosys)^53^ and searched against the UniProtKB/Swiss-Prot database (retrieved: 21.10.2021, 20387 entries) as well as a database containing protein sequences longer than 7 amino acids predicted from translation of uORFs. Search parameters were set to non-specific digestion and a peptide length of 7-15 amino acids. Carbamidomethylation of cysteine and oxidation of methionine were included as variable modifications. Results were reported with 1% FDR at the peptide level. Peptides identified by Spectronaut were further analyzed using NetMHCpan 4.1 binding predictions^54^, Gibbs 2.0 clustering of peptide sequences^55^, and retention time prediction by DeepLC^56^. uORF/uoORF-derived peptide sequences were manually validated using Skyline (version 22)^57^ by comparison against spectral libraries *in silico* predicted using PROSIT^58^. Normalized spectral angles (NSAs) were calculated as described previously^59^. Quantification of HLA class I-presented peptides was performed as described previously^60^ using the raw output at the MS2 level from Spectronaut 17.6 with cross-run normalization disabled and a custom script in the R programming language. Peptides with an FDR ≤ 0.05 and an foldchange > 2 were defined as “hits” while peptides with an FDR ≤ 0.2 and an foldchange ≥ 1.5 were defined as “candidates”. All results were visualized using in-house developed R scripts

### Mice

OT-I animales were bred in our animal facility. All mice were maintained in a pathogen-free facility and used according to the German Cancer Research Center and following permission by the controlling government office (Regierungspräsidium Karlsruhe) according to the German Animal Protection Law, and in compliance with the EU Directive on animal welfare, Directive 2010/63/EU.

### CD8+ T cell isolation and culture

Primary naive CD8+ T OT-I cells were isolated using the MojoSort Mouse CD8 T cell isolation kit (BioLegend, 480007) and subsequently activated for 72 hours on plates coated with 2 μg/ml anti-CD3 (BioXCell, BE0001-1) and 2 μg/ml anti-CD28 (BioXCell, BE0015-1) at 37°C. The T cells were maintained in RPMI 1640 (Thermo Fisher Scientific, 21875-034) supplemented with 10% FBS, 1% penicillin-streptomycin, 1 mM sodium pyruvate (Thermo Fisher Scientific, 11360-070), 20 mM HEPES (Sigma, H4034), 50 µM β-mercaptoethanol (Sigma, M6250), and 10 ng/ml murine IL-2 (BioLegend, 575404).

### Quantitative real-time PCR

A total of 500 ng of RNA was reverse-transcribed using LunaScript RT Supermix (NEB, M3010L). Quantitative real-time PCR was subsequently performed with Luna Universal qPCR Mix (NEB, M3003X). Ct values were obtained using the QuantStudio 5 RT qPCR System and analyzed with QuantStudio Design and Analysis Software v2.6.0. mRNA fold change of target genes was calculated using the ΔΔCt method, with mRNA expression normalized to GAPDH. The primers used in this study are listed in Supplementary Dataset 11.

### 5’UTR cloning

A synthetic DNA template containing the 5’ UTR and the coding sequence for the SIINKEKL peptide was amplified by PCR using primers with overlapping sequences complementary to the pGL3-Promoter vector (Promega). Following PCR, the fragments were gel-purified to remove any non-specific products. The pGL3-Promoter vector was linearized using HindIII-HF (NEB, catalog no. R3104). The purified insert and linearized vector were then combined with NEBuilder HiFi DNA Assembly Master Mix (NEB, catalog no. E2621) and incubated at 50°C for 60 minutes. The assembled product was transformed into competent cells for propagation and further analysis. Primers and synthetic DNA templates used in this study are listed in Supplementary Dataset 11.

### T cell killing assay

CD8^+^ OT-I T cells were co-cultured with mouse cancer cells that express the specified uORF- SIINFEKL reporters at a ratio of 1:2. Following 24 hours of incubation at 37°C in a 5% CO_2_ atmosphere, the cells were washed with PBS and stained with crystal violet to evaluate the killing efficiency. Imaging was conducted using the Dual Lens System V850 Pro Scanner (Epson), and colony area was quantified using a previously published ImageJ plugin^61^.

### Flow cytometry

TC1 cells were transfected with the uORF-SIINFEKL reporters using Lipofectamine 3000 (Thermo Fisher). 24 hours post-transfection, cells were synchronized in mitosis by treatment with 1 µM Taxol (Santa Cruz, sc-201439) for an additional 16 hours. Following mitotic arrest, cells were washed with PBS, detached using PBS-EDTA, and then pelleted. The cells were subsequently washed with PBS containing 0.5% BSA and incubated on ice and in the dark with APC-conjugated anti-mouse H-2Kb-SIINFEKL antibodies (Biolegend, clone 25-D1.16, #141606; 1:200) for 30 minutes. After incubation, the cells were washed twice with PBS containing 0.1% BSA and analyzed using a FACS Canto II cytometer (Thermo Fisher). Data analysis was conducted with FlowJo V10.4 software (FlowJo).

### IFN-γ quantification

Cytokine release from CD8+ T cells was measured from the cell supernatant using the ELISA MAX Deluxe Set Mouse IFN-γ (BioLegend, 430815), following the manufacturer’s guidelines. Each sample was analyzed with the Multiskan FC plate reader (Thermo Fisher Scientific), using absorbance readings at 450 nm and 570 nm for subtraction. Final concentrations were calculated using a 4-parameter logistic curve-fitting algorithm in GraphPad Prism.

## Data availability

The sequence data from this study have been submitted to the GEO repository: GSE281253 (https://www.ncbi.nlm.nih.gov/geo/query/acc.cgi?acc=GSE281253, token: cxqrkygatnkfdup).

The mass spectrometry proteomics data have been deposited to the ProteomeXchange Consortium via the PRIDE partner repository with the dataset identifier PXD057839. Reviewers can access the data with the username and password “krYPSAic53Ox”.

## Acknowledgements

We thank Wilhelm Palm for advice and critical discussions. We thank Chong Sun for sharing reagents and technical advice. This work was funded in part by grants of the European Research Council “DualRP” (ERC StG No. 759579) and the German Research Foundation (DFG 504774163 and DFG 545215964) to F.L.-P. R.A. is supported by the Dutch Cancer Society (KWF-13647), the European Research Council (Horizon ERC-2023-ADG-101141245), the Dutch Science Organization (OCENW.M.22.001-16221), and the AvL Foundation. A.K. and R.DP. are supported by fellowships of the Helmholtz International Graduate School. Z.T. is supported by a scholarship from the China Scholarship Council (CSC).

## Author contributions

A.K., R.A., and F.L.-P. conceived the project, designed all the experiments, and wrote the manuscript. Methodology and data acquisition: A.K., J.P.B., R.DP., Z.T., F.M.T., H.S. Manuscript revision: A.K., A.B.R., R.A., and F.L-P.

## References

1. Chan, K.-S., Koh, C.-G. & Li, H.-Y. Mitosis-targeted anti-cancer therapies: where they stand. Cell Death Dis. 3, e411 (2012).

2. Tanenbaum, M. E., Stern-Ginossar, N., Weissman, J. S. & Vale, R. D. Regulation of mRNA translation during mitosis. Elife 4, (2015).

3. Stumpf, C. R., Moreno, M. V., Olshen, A. B., Taylor, B. S. & Ruggero, D. The Translational Landscape of the Mammalian Cell Cycle. Mol. Cell 52, 574–582 (2013).

4. Park, J.-E., Yi, H., Kim, Y., Chang, H. & Kim, V. N. Regulation of Poly(A) Tail and Translation during the Somatic Cell Cycle. Mol. Cell 62, 462–471 (2016).

5. Shuda, M. et al. CDK1 substitutes for mTOR kinase to activate mitotic cap-dependent protein translation. Proc. Natl. Acad. Sci. U. S. A. 112, 5875–5882 (2015).

6. Haneke, K. et al. CDK1 couples proliferation with protein synthesis. J. Cell Biol. 219, (2020).

7. Imami, K. et al. Phosphorylation of the ribosomal protein RPL12/uL11 affects translation during mitosis. Mol. Cell 72, 84–98.e9 (2018).

8. Jordan, M. A. & Wilson, L. Microtubules as a target for anticancer drugs. Nat. Rev. Cancer 4, 253–265 (2004).

9. Wani, M. C., Taylor, H. L., Wall, M. E., Coggon, P. & McPhail, A. T. Plant antitumor agents. VI. The isolation and structure of taxol, a novel antileukemic and antitumor agent from Taxus brevifolia. J. Am. Chem. Soc. 93, 2325–2327 (1971).

10. Rowinsky, E. K. & Donehower, R. C. Paclitaxel (taxol). N. Engl. J. Med. 332, 1004–1014 (1995).

11. Zhu, L. & Chen, L. Progress in research on paclitaxel and tumor immunotherapy. Cell. Mol. Biol. Lett. 24, 40 (2019).

12. Alalawy, A. I. Key genes and molecular mechanisms related to Paclitaxel Resistance. Cancer Cell Int. 24, 244 (2024).

13. Schiff, P. B. & Horwitz, S. B. Taxol stabilizes microtubules in mouse fibroblast cells. Proc. Natl. Acad. Sci. U. S. A. 77, 1561–1565 (1980).

14. Rowinsky, E. K., Donehower, R. C., Jones, R. J. & Tucker, R. W. Microtubule changes and cytotoxicity in leukemic cell lines treated with taxol. Cancer Res. 48, 4093–4100 (1988).

15. Dumontet, C. & Jordan, M. A. Microtubule-binding agents: a dynamic field of cancer therapeutics. Nat. Rev. Drug Discov. 9, 790–803 (2010).

16. Ruiz Cuevas, M. V., et al. Most non-canonical proteins uniquely populate the proteome or immunopeptidome. Cell Rep. 34, 108815 (2021).

17. Chong, C. et al. Integrated proteogenomic deep sequencing and analytics accurately identify non-canonical peptides in tumor immunopeptidomes. Nat. Commun. 11, 1293 (2020).

18. 18. Ji, Z., Song, R., Regev, A. & Struhl, K. Many lncRNAs, 5’UTRs, and pseudogenes are translated and some are likely to express functional proteins. Elife 4, (2015).

19. Prensner, J. R. et al. Noncanonical open reading frames encode functional proteins essential for cancer cell survival. Nat. Biotechnol. 39, 697–704 (2021).

20. Vieira de Souza, E., et al. Rp3: Ribosome profiling-assisted proteogenomics improves coverage and confidence during microprotein discovery. Nat. Commun. 15, 6839 (2024).

21. van Heesch, S. et al. The translational landscape of the human heart. Cell 178, 242–260.e29 (2019).

22. Chen, J. et al. Pervasive functional translation of noncanonical human open reading frames. Science 367, 1140–1146 (2020).

23. Chothani, S. P. et al. A high-resolution map of human RNA translation. Mol. Cell 82, 2885–2899.e8 (2022).

24. Ferreira, H. J. et al. Immunopeptidomics-based identification of naturally presented non-canonical circRNA-derived peptides. Nat. Commun. 15, 2357 (2024).

25. Ouspenskaia, T. et al. Unannotated proteins expand the MHC-I-restricted immunopeptidome in cancer. Nat. Biotechnol. 40, 209–217 (2022).

26. Anderson, D. M. et al. A micropeptide encoded by a putative long noncoding RNA regulates muscle performance. Cell 160, 595–606 (2015).

27. Ho, L. et al. ELABELA is an endogenous growth factor that sustains hESC self-renewal via the PI3K/AKT pathway. Cell Stem Cell 17, 635 (2015).

28. Duffy, E. E. et al. Developmental dynamics of RNA translation in the human brain. Nat. Neurosci. 25, 1353–1365 (2022).

29. Hurwitz, B. et al. The integrated stress response remodels the microtubule-organizing center to clear unfolded proteins following proteotoxic stress. Elife 11, (2022).

30. Sendoel, A. et al. Translation from unconventional 5’ start sites drives tumour initiation. Nature 541, 494–499 (2017).

31. Velásquez, C. et al. Mitotic protein kinase CDK1 phosphorylation of mRNA translation regulator 4E-BP1 Ser83 may contribute to cell transformation. Proc. Natl. Acad. Sci. U. S. A. 113, 8466–8471 (2016).

32. Ingolia, N. T., Lareau, L. F. & Weissman, J. S. Ribosome profiling of mouse embryonic stem cells reveals the complexity and dynamics of mammalian proteomes. Cell 147, 789– 802 (2011).

33. Lee, S. et al. Global mapping of translation initiation sites in mammalian cells at single- nucleotide resolution. Proc. Natl. Acad. Sci. U. S. A. 109, E2424–32 (2012).

34. Erhard, F. et al. Improved Ribo-seq enables identification of cryptic translation events. Nat. Methods 15, 363–366 (2018).

35. Jürgens, L. & Wethmar, K. The emerging role of uORF-encoded uPeptides and HLA uLigands in cellular and tumor biology. Cancers (Basel) 14, 6031 (2022).

36. Schmid, P. et al. Atezolizumab and nab-paclitaxel in advanced triple-negative breast cancer. N. Engl. J. Med. 379, 2108–2121 (2018).

37. Marash, L. et al. DAP5 promotes cap-independent translation of Bcl-2 and CDK1 to facilitate cell survival during mitosis. Mol. Cell 30, 447–459 (2008).

38. Olan, I. et al. HMGA1 orchestrates chromatin compartmentalization and sequesters genes into 3D networks coordinating senescence heterogeneity. Nat. Commun. 15, 6891 (2024).

39. Sgubin, M. et al. HMGA1 positively regulates the microtubule-destabilizing protein stathmin promoting motility in TNBC cells and decreasing tumour sensitivity to paclitaxel. Cell Death Dis. 13, 429 (2022).

40. Dersh, D., Yewdell, J. W. & Wei, J. A SIINFEKL-based system to measure MHC class I antigen presentation efficiency and kinetics. Methods Mol. Biol. 1988, 109–122 (2019).

41. Chen, Y. et al. Nab-paclitaxel promotes the cancer-immunity cycle as a potential immunomodulator. Am. J. Cancer Res. 11, 3445–3460 (2021).

42. Yang, Q. et al. Nanomicelle protects the immune activation effects of Paclitaxel and sensitizes tumors to anti-PD-1 Immunotherapy. Theranostics 10, 8382–8399 (2020).

43. Bracci, L., Schiavoni, G., Sistigu, A. & Belardelli, F. Immune-based mechanisms of cytotoxic chemotherapy: implications for the design of novel and rationale-based combined treatments against cancer. Cell Death Differ. 21, 15–25 (2014).

44. Ly, J. et al. Nuclear release of eIF1 restricts start-codon selection during mitosis. Nature (2024) doi:10.1038/s41586-024-08088-3.

45. Mudge, J. M. et al. Standardized annotation of translated open reading frames. Nat. Biotechnol. 40, 994–999 (2022).

46. Ji, Z. RibORF: Identifying genome-wide translated open reading frames using ribosome profiling. Curr. Protoc. Mol. Biol. 124, e67 (2018).

47. Calviello, L., Hirsekorn, A. & Ohler, U. Quantification of translation uncovers the functions of the alternative transcriptome. Nat. Struct. Mol. Biol. 27, 717–725 (2020).

48. Choudhary, S., Li, W. & D Smith, A. Accurate detection of short and long active ORFs using Ribo-seq data. Bioinformatics 36, 2053–2059 (2020).

49. Prensner, J. R. et al. What can Ribo-Seq, immunopeptidomics, and proteomics tell us about the noncanonical proteome? Mol. Cell. Proteomics 22, 100631 (2023).

50. Goyal, A. et al. DNMT and HDAC inhibition induces immunogenic neoantigens from human endogenous retroviral element-derived transcripts. Nat. Commun. 14, 6731 (2023).

51. Stopfer, L. E., Mesfin, J. M., Joughin, B. A., Lauffenburger, D. A. & White, F. M. Multiplexed relative and absolute quantitative immunopeptidomics reveals MHC I repertoire alterations induced by CDK4/6 inhibition. Nat. Commun. 11, 2760 (2020).

52. Chong, C. et al. High-throughput and sensitive immunopeptidomics platform reveals profound interferonγ-mediated remodeling of the human leukocyte antigen (HLA) ligandome. Mol. Cell. Proteomics 17, 533–548 (2018).

53. Bruderer, R. et al. Extending the limits of quantitative proteome profiling with data-independent acquisition and application to acetaminophen-treated three-dimensional liver microtissues. Mol. Cell. Proteomics 14, 1400–1410 (2015).

54. Reynisson, B., Alvarez, B., Paul, S., Peters, B. & Nielsen, M. NetMHCpan-4.1 and NetMHCIIpan-4.0: improved predictions of MHC antigen presentation by concurrent motif deconvolution and integration of MS MHC eluted ligand data. Nucleic Acids Res. 48, W449–W454 (2020).

55. Andreatta, M., Alvarez, B. & Nielsen, M. GibbsCluster: unsupervised clustering and alignment of peptide sequences. Nucleic Acids Res. 45, W458–W463 (2017).

56. Bouwmeester, R., Gabriels, R., Hulstaert, N., Martens, L. & Degroeve, S. DeepLC can predict retention times for peptides that carry as-yet unseen modifications. Nat. Methods 18, 1363–1369 (2021).

57. Pino, L. K. et al. The Skyline ecosystem: Informatics for quantitative mass spectrometry proteomics. Mass Spectrom. Rev. 39, 229–244 (2020).

58. Gessulat, S. et al. Prosit: proteome-wide prediction of peptide tandem mass spectra by deep learning. Nat. Methods 16, 509–518 (2019).

59. Toprak, U. H. et al. Conserved peptide fragmentation as a benchmarking tool for mass spectrometers and a discriminating feature for targeted proteomics. Mol. Cell. Proteomics 13, 2056–2071 (2014).

60. Becker, J. P. et al. NMD inhibition by 5-azacytidine augments presentation of immunogenic frameshift-derived neoepitopes. iScience 24, 102389 (2021).

61. Guzmán, C., Bagga, M., Kaur, A., Westermarck, J. & Abankwa, D. ColonyArea: an ImageJ plugin to automatically quantify colony formation in clonogenic assays. PLoS One 9, e92444 (2014).

