## Supplementary information for "Upstream open reading frame translation enhances immunogenic peptide presentation in mitotically arrested cancer cells"

#### Supplementary Figures 1 to 5

### Supplementary Figure 1

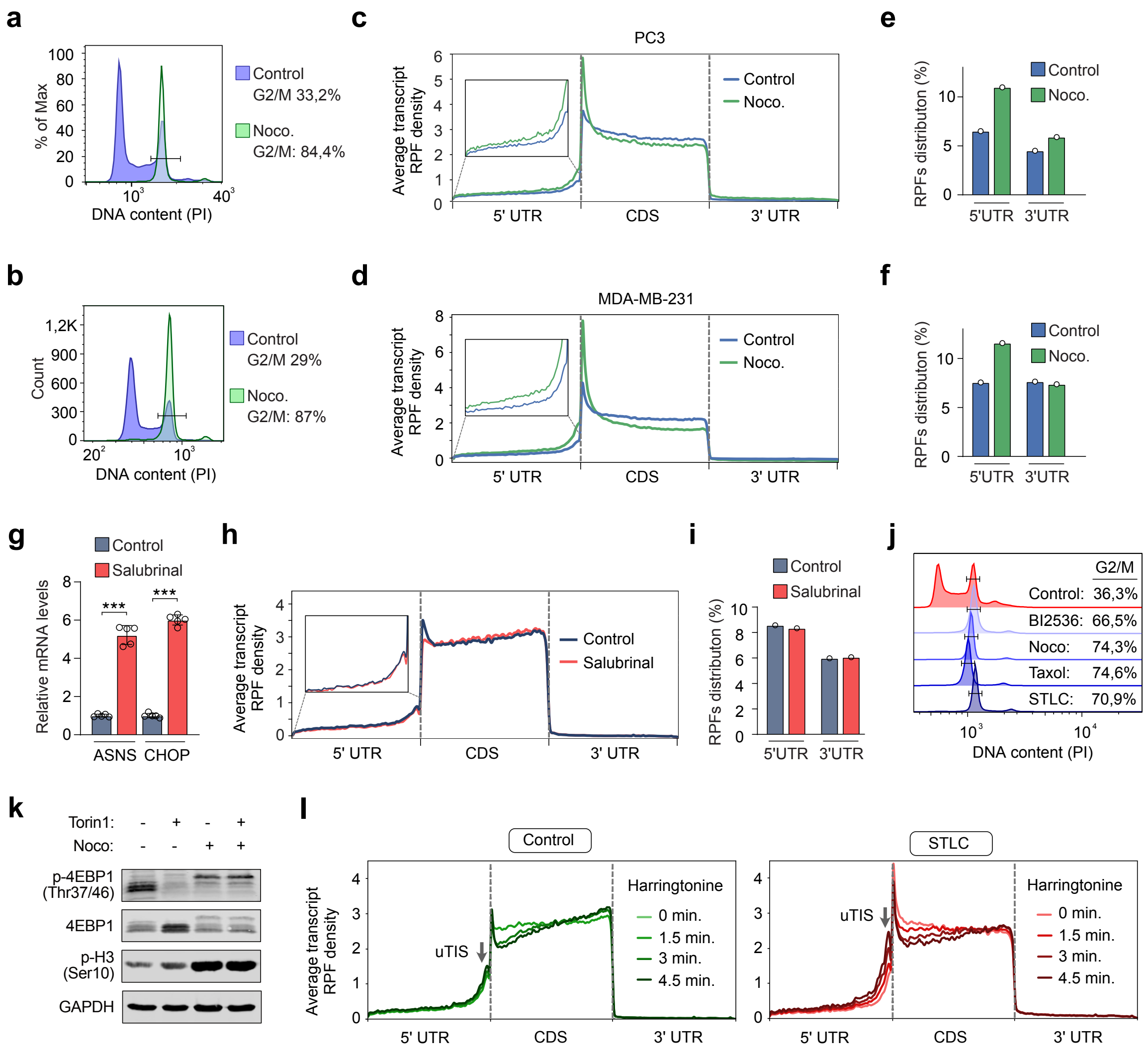

**Supplementary Figure 1. Impact of mitotic arrest agents on RPF distribution in cancer cells.**

**a-b,** Representative propidium iodide stainings of PC3 (a) and MDA-MB-231 (b) cells arrested in mitosis with Nocodazole (Noco. 0,5 μM) for 16 hours.

**c-d,** Metagene profiles of RPFs in proliferating and mitotically arrested PC3 (c) and MDA-MB-231 (d) cells treated with Nocodazole (0.5 μM; 16 hours).

**e-f,** Quantification of RPF distribution in the 5' UTR of PC3 (e) and MDA-MB-231 (f) cells treated with vehicle (Control) or Nocodazole (0,5 μM) for 16 hours.

**g,** qRT-PCR analysis of the indicated genes in U2OS cells treated with vehicle (Control) or Salubrinal (50 μM; 16 hours). Data represent mean ± SD from biologically independent experiments ($n$ = 5). $p$-values were calculated using a two-tailed unpaired $t$-test. ***$P < 0.001$.

**h,** Metagene profiles of RPFs in U2OS cells treated with vehicle (Control) or Salubrinal (50 μM; 16 hours).

**i**, Quantification of RPF distribution in the 5' UTR and 3'UTR of U2OS cells treated with vehicle (Control) or Salubrinal (50 μM; 16 hours).

**j,** Representative propidium iodide stainings of U2OS cells treated with vehicle (Control) or BI2536. (0,1 μM), Nocodazole (0,5 μM), Taxol (1 μM), or STLC (1 μM) for 16 hours.

**k,** Immunoblot of phospho-4E-BP1 (p-4E-BP1, Thr37/46) and total 4E-BP1 in U2OS cells treated with vehicle (Control), Nocodazole (0,5 μM), or Nocodazole (0,5 μM) + Torin1 (250 nM). Cells were treated with Nocodazole for 16 hours and with Torin1 for 2 hours.

**l**, Metagene profiles of RPFs in MDA-MB-231 cells treated with vehicle (Control) or STLC (5 μM) for 16 hours. Cells were harvested as described in Fig. 1i. uTIS, upstream translation initiation site.

#### Supplementary Figure 2

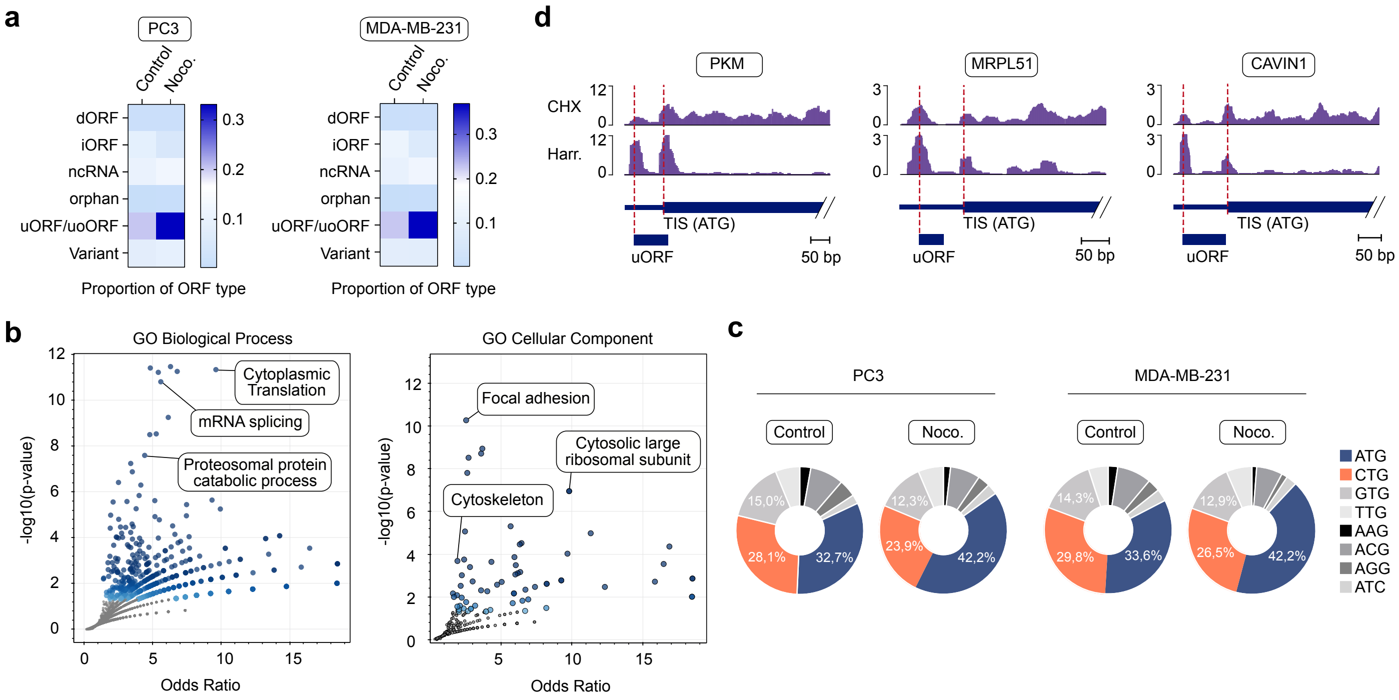

**Supplementary Figure 2. Analysis of uORF/uoORF Translation in mitotically arrested Cancer Cells**

**a,** Proportion of ORF categories described in Fig. 2a in PC3 (left panel) and MDA-MB-231 cells (right panel) treated with vehicle (Control) or Nocodazole (0.5 μM) for 16 hours.

**b**, Volcano plots showing enriched terms from the Gene Ontology (GO) Biological Process (left panel) and Cellular Component (right panel) for the common genes described in Fig. 2c. Each point represents a term, with the x-axis showing the odds ratio and the y-axis showing the -log10(p-value). Larger and darker points signify terms with higher enrichment significance in the input gene set.

**c,** Percentage of uORF/uoORF translation initiation sites in PC3 (left panel) and MDA-MB-231 (right panel) cells treated with vehicle (Control) or Nocodazole (0.5 μM) for 16 hours.

**d,** Read distribution from translation initiation site sequencing of representative uORFs/uoORFs in U2OS cells arrested in mitosis (Nocodazole, 0.5 μM, 16 hours). CHX, Cycloheximide; Harr., Harringtonine.

#### Supplementary Figure 3

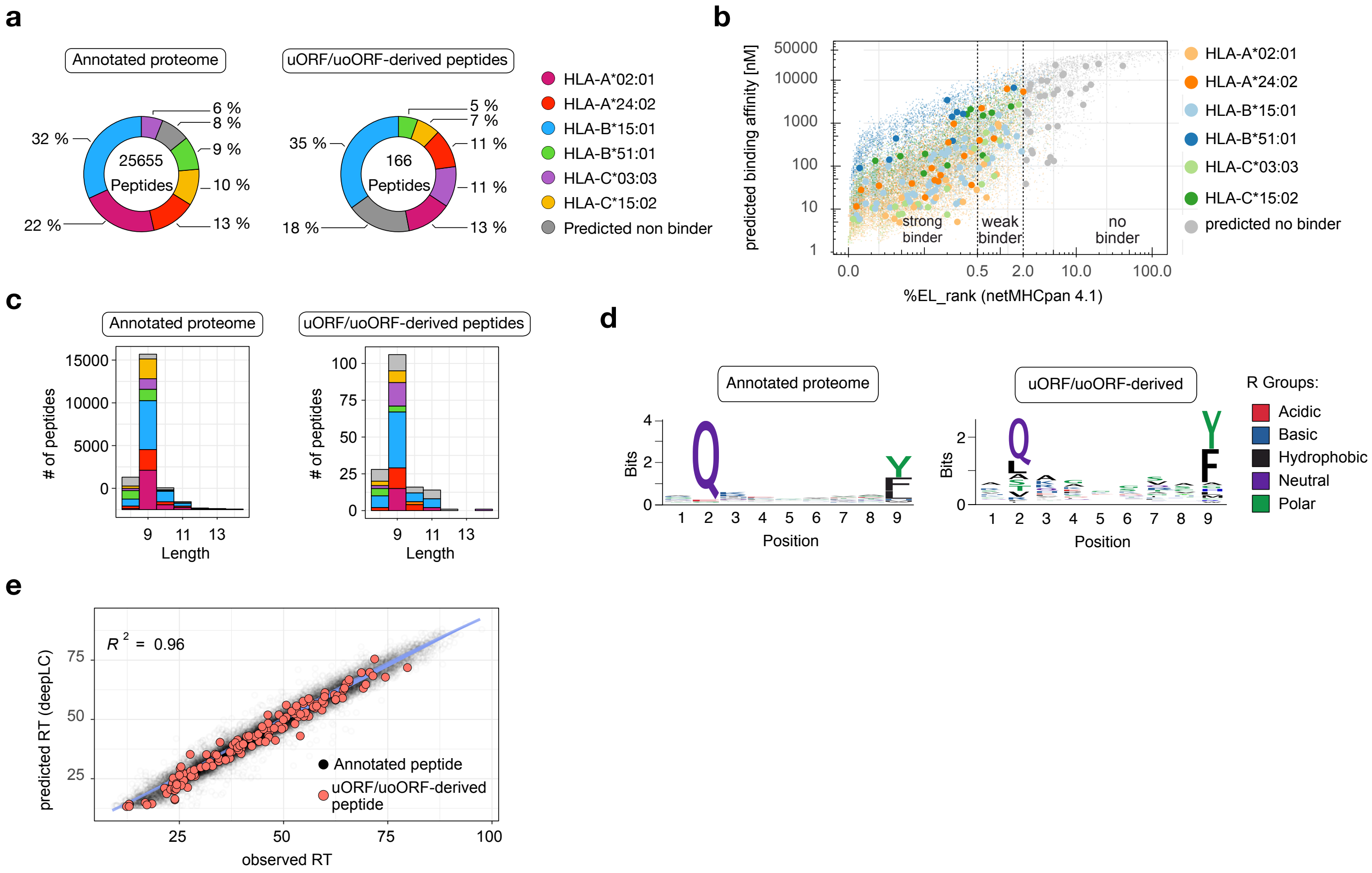

**Supplementary Figure 3. Biochemical characteristics of annotated and uORF/uoORF-derived HLA-presented peptides in SUM-159PT cells.**

**a,** Quantification of peptides in the annotated proteome (left panel) and uORF/uoORF-derived proteome (right panel) in proliferating and mitotically arrested SUM-159PT cells. The distribution of predicted binding to HLA alleles in SUM-159PT cells is presented.

**b,** Percentage of eluted ligand (EL) peptides predicted by NetMHCpan-4.1 plotted against predicted binding affinity, for peptides from the annotated proteome (small dots) and uORF/uoORF-derived peptides (large dots) of proliferating and mitotically arrested SUM-159PT cells. Predicted binding to HLA alleles in SUM-159PT cells is displayed. Peptides are classified as strong binders (%EL rank 0–0.5), weak binders (%EL rank 0.5–2), or non-binders (%EL rank 2–100).

**c,** Length distribution of detected peptides from the annotated proteome (left panel) and uORF/uoORF-derived peptides (right panel) in proliferating and mitotically arrested SUM-159PT cells. The proportion of predicted binding to HLA alleles in SUM-159PT cells is shown.

**d.** Peptide motif plots for unique peptides from the annotated proteome (4,402) and unique peptides derived from uORFs/uoORFs (59), confidently identified as binding to the SUM-159PT allele HLA-B*15:01.

**e,** Observed retention time (RT) indices plotted against predicted RT indices for peptides from the annotated (black) and uORF/uoORF-derived (red) proteomes, across all HLA alleles in proliferating and mitotically arrested SUM-159PT cells. $R^2$ indicates the Pearson correlation coefficient.

#### Supplementary Figure 4

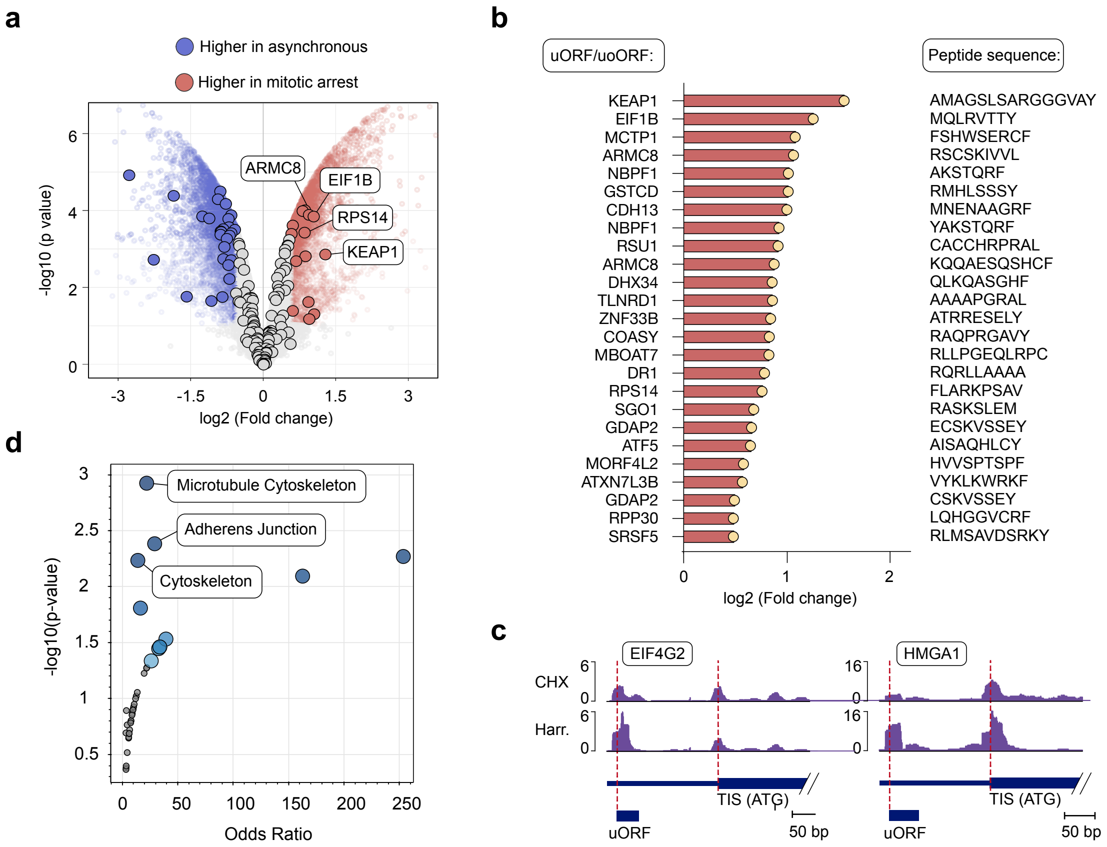

**Supplementary Figure 4. Label-Free quantification of HLA-presented uORF/uoORF-derived peptides in mitotically arrested cancer cells.**

**a,** Volcano plot showing label-free quantification of the immunopeptidome in Taxol-treated versus DMSO-treated SUM-159PT cells, highlighting the genes expressing uORF/uoORF-derived peptides. Peptides with a log2 fold change greater than 0.5 and adjusted p-value less than 0.05 are in red, while those with a log2 fold change below -0.5 and adjusted p-value less than 0.05 are in blue. The analysis was conducted using an empirical Bayes moderated *t*-test with two-sided *p*-values.

**b,** Mitotic arrest-induced uORF/uoORF-derived peptides in SUM-159PT cells, listing host gene name, log2 fold change (FC) of peptide abundance, and peptide sequences.

**c,** RPF reads distribution from translation initiation site sequencing of representative uORFs/uoORFs in U2OS cells arrested in mitosis. CHX, Cycloheximide. Harr, Harringtonine.

**d,** Gene Ontology (GO) Biological Process terms for the upregulated peptides in Fig. 4a, with each point representing a specific term. The x-axis displays the odds ratio, while the y-axis shows the -log10(*p*-value). Larger, darker points indicate terms with greater enrichment significance in the input gene set.

### Supplementary Figure 5

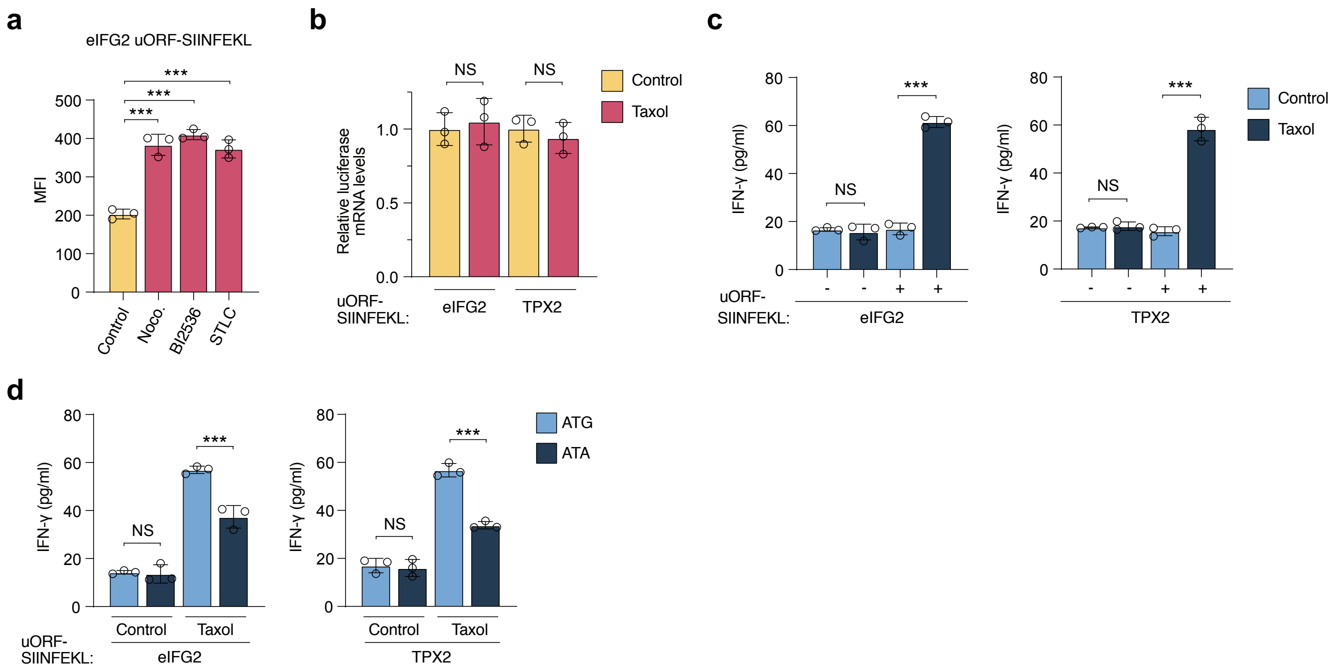

**Supplementary Figure 5. Effect of mitotic arrest on peptide presentation and IFNγ production in TC1 Cells.**

**a,** Detection of the p:MHC complex SIINFEKL:H-2K$^b$ by flow cytometry, shown as Median fluorescence intensity (MFI), in TC1 cells transfected with the indicated reporters. Cells were treated with vehicle (Control) or BI2536 (0.1 µM), Nocodazole (0.5 µM), or STLC (5 µM) for 16 hours. Data represent mean ± SD from biologically independent experiments (*n* = 3). *p*-values were calculated using a two-tailed unpaired *t*-test. ***$P < 0.001$.

**b,** qRT-PCR of firefly luciferase in TC1 cells transfected with the indicated reporters. Cells were treated with vehicle (Control) or Taxol (1 µM) for 16 hours. Data represent mean ± SD from biologically independent experiments (*n* = 5). *p*-values were calculated using a two-tailed unpaired *t*-test. ***$P < 0.001$.

**c-d**, Quantification of IFN-γ levels produced by CD8+ OT-I T cells when co-cultured with TC1 cells. TC1 cells were transfected with the indicated uORF-SIINFEKL reporters and arrested in mitosis with Taxol (1 µM) for 16 hours. Control cells were treated with DMSO for the same period. Data represent mean ± SD from biologically independent experiments (*n* = 3). *p*-values were calculated using a two-tailed unpaired *t*-test. ***$P < 0.001$.
